# Spatiotemporal evidence accumulation through saccadic sampling for object recognition

**DOI:** 10.1101/2024.09.05.611201

**Authors:** Zhihao Zheng, Jiaqi Hu, Gouki Okazawa

## Abstract

Visual object recognition has been extensively studied under fixation conditions, but our natural viewing involves frequent saccadic eye movements that scan multiple local informative features within an object (e.g., eyes and mouth in a face image). These saccades would contribute to object recognition by subserving the integration of sensory information across local features, but mechanistic models that explain human behavior during natural object recognition have yet to be established due to the presumed complexity of the interactions between the visual and oculomotor systems. Here, we employ a framework of perceptual decision making and show that human face and object categorization behavior with saccades can be quantitatively explained by a model that simply accumulates the sensory evidence available at each moment. Our model can fit human object-recognition behavior even under conditions in which people freely make saccades to scan local features, departing from past studies that required controlled eye movements to test trans-saccadic integration. Moreover, further experimental results confirmed that active saccade commands (efference copy) do not substantially contribute to behavioral performance. Therefore, we propose that object recognition with saccades can be approximated by a parsimonious decisionmaking model without assuming complex interactions between the visual and oculomotor systems.

## Introduction

Object recognition is often studied as the process of extracting object information from a static image, but during a natural visual experience, our visual system is bombarded with frequent and drastic changes in the retinal image due to our own eye movements. Saccadic eye movements, which occur on average 1-3 times per second (1), help us search for and focus on an important object in a visual scene by shifting the center of gaze from one object to another (2). However, even when looking at a single object, we often make saccades within the object to scan multiple local features. For example, in classic demonstrations of human eye tracking by A. Yarbus (1967), people viewing a face made frequent saccades across key facial features such as eyes and mouth.

A number of studies have investigated eye movement patterns during object viewing (4–6) and have suggested the importance of saccades in face and object recognition (5–7). People fixate on the most informative region of an image during face recognition (8, 9) and adopt different saccade patterns depending on task demands (5, 6, 10). Limiting saccades or the number of fixations impairs object recognition and learning performance (11–13). Integration of visual information across saccades (i.e., trans-saccadic integration) has been well demonstrated in studies using simplified stimuli such as Gabor orientations (14, 15), color patches (16), or 2D shapes (17–20). These studies have shown that humans integrate the same visual feature viewed at the fovea and periphery across a saccade (21).

However, important questions remain to be addressed regarding the mechanisms of how eye movements contribute to object vision. First, most existing work on trans-saccadic integration has used simple visual features (such as Gabor) (14–16, 22), whereas the recognition of complex object images (e.g., faces) may pose different challenges because saccades bring different complex features (e.g., eyes and mouth) to both the fovea and periphery. Existing models of form and object vision suggest that sensory evidence is integrated even in such cases (23–25), but an empirical test with natural images would be crucial to verify this. Second, the integration during saccades may require additional neural processes, such as combining visual inputs with efference copy from the oculomotor system (26, 27); whether such processes play a major role in object recognition with saccades should also be tested empirically. Finally, trans-saccadic integration is usually studied under conditions in which participants are explicitly asked to make a saccade in a controlled manner (14–20). It remains to be seen whether people integrate evidence even when they make free saccades during object viewing.

Here, we develop a quantitative framework for measuring and modeling object recognition behavior with saccades to address all these questions. Our key innovation is to employ a theory of perceptual decision making (28) and study object recognition behavior as a process of accumulating sensory evidence over time to commit to a choice (29–32). Using behavioral paradigms with parametrically controlled object and face stimuli, we demonstrate that a simple model that accumulates evidence of local visual features across saccades is sufficient to quantitatively account for participants’ object categorization behavior even when they view an image freely. Moreover, further experiments confirmed that participants’ behavioral performance was minimally affected by the efference copy of oculomotor signals. We conclude that object recognition with saccades can be approximated by a parsimonious model that accumulates available sensory evidence from dynamic retinal images without assuming complex interactions between the visual and oculomotor systems under controlled conditions in which trained participants repeatedly categorize objects.

## Results

### Saccadic sampling of local features during object recognition

We designed an object categorization task in which participants freely made eye movements inside a stimulus to report its category. We assigned participants to perform either facial identity categorization, facial expression categorization, or car categorization (Fig. 1A) and aimed to reveal the behavioral patterns common to these three conditions. A stimulus in each trial was sampled from a morph continuum of two prototype images (e.g., two facial identities in the identity categorization, which corresponded to -100 and 100% morph; Fig. 1A and Supplementary Fig. 1A), and the participants were asked to report which prototype category the stimulus was closer to. Before stimulus onset, participants were required to look at a fixation point whose position was randomly selected from six possible peripheral locations (8 - 11.5° away from the monitor center; Fig. 1B). Following stimulus onset, the participants had to immediately make a saccade to the stimulus and could then look at any part inside it. They subsequently reported their decisions by pressing a button as soon as they were ready (reaction-time task; Fig. 1B). Reaction times (RTs) were defined as the time between the fixation on the stimulus and the button press.

**Fig 1.**
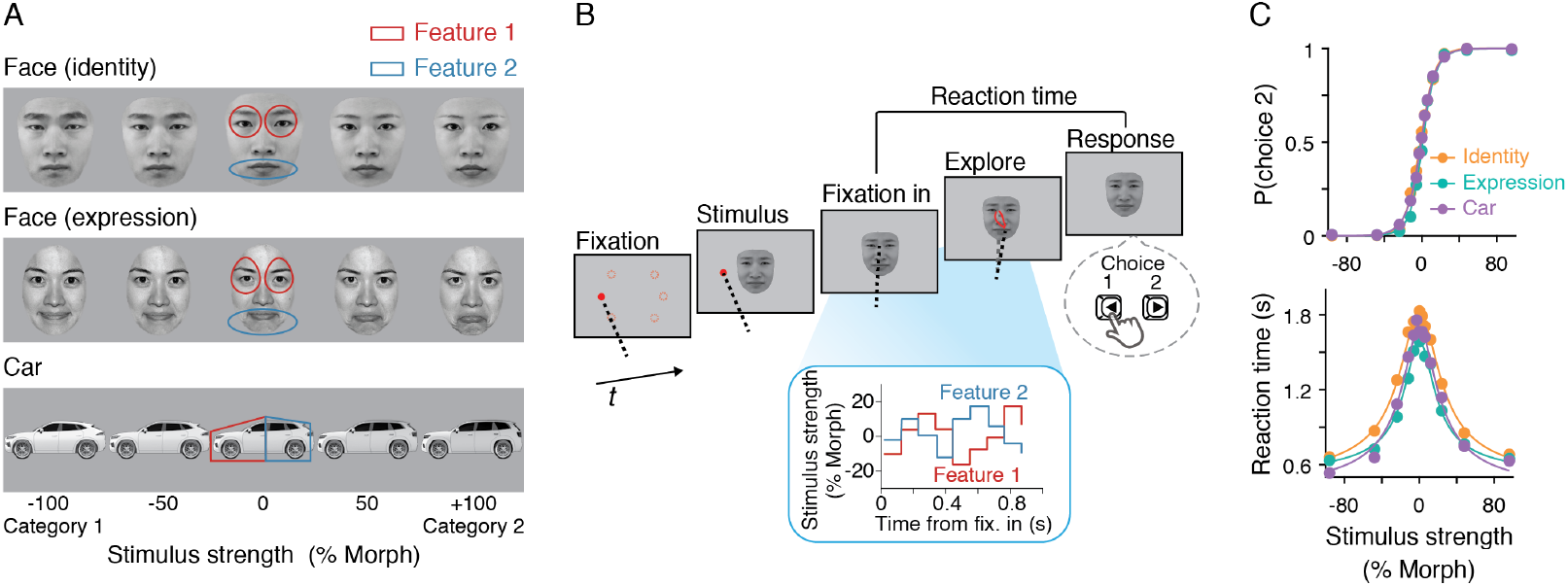
Object categorization task with free saccades. (**A**) Participants were assigned to perform either facial identity, expression, or car categorization (*n* = 9). In each trial, participants viewed a stimulus chosen from a morph continuum ranging from -100 to 100% morph levels between two prototype images. Participants were then required to report which prototype the stimulus was closer to. We defined two informative features (red and blue contour lines) and morphed images inside these two regions. The face images were from the Nim Face set (33) and the Tsinghua Facial Expression Database (34) and presented with permission. The same face images were used in the subsequent figures. (**B**) Participants began each trial by fixating on a red dot that appeared at one of six possible locations on the screen. Shortly afterward, a stimulus appeared, and participants were required to make a saccade to the stimulus. Thereafter, participants could view any part of the image until they reported the stimulus category by pressing one of two buttons as soon as they were ready (reaction-time task). During stimulus viewing, the morph levels of the two informative features fluctuated randomly every 106.7 ms, while their mean was maintained constant within a trial (SD: 20%). This fluctuation allowed us to examine the weighting of each feature during decision making. An example movie of the dynamic face stimuli can be found in Okazawa et al. (2021). (**C**) Participants showed stereotypical psychometric and chronometric curves as a function of the mean morph levels. Lines represent logistic and hyperbolic tangent fits for psychometric and chronometric functions, respectively (Eq. 1 and 2). Plots for each participant are shown in Supplementary Fig. 2.

To assess how participants sampled the local features during decision making, we defined two informative features for each stimulus set (eyes and mouth in the face sets; front and rear parts in the car set; Fig. 1A) and added random fluctuations to their morph levels every 106.7 ms (eight monitor frames) during stimulus presentation (Fig. 1B inset; fluctuation SD, 20% morph). The mean morph levels of the two features were maintained identical and constant within each trial. The morph level outside of the two features was always set to 0% and remained uninformative. This design allowed us to use psychophysical reverse correlation (29, 30, 35) and test how each feature at each moment during stimulus viewing influenced the participants’ decisions. Two features were sufficiently separated for saccades to be made between them (inter-feature distance, 5°), and at the same time, the image sizes were within the range of naturalistic viewing conditions (36).

We first confirmed that the participants showed stereotypical choice accuracy and RTs under the three stimulus conditions (Fig. 1C). Hereafter, we focus on the results qualitatively consistent across the three conditions and present the results either individually or averaged across conditions depending on the purpose of visualization (where averaged results are presented, individual results are shown in the Supplementary Figures). In all conditions, choice accuracy was monotonically modulated by the morph level (logistic regression slope *α*_1_ = 13.7 ± 1.2, mean ± S.E.M. across participants, Eq. 1; *t*_(8)_ = 11, *p* = 4.0 ×10^−6^, two-tailed *t*-test). RTs were systematically longer for lower morph lev-els (*β*_1_ = 7.14 ± 0.52 fitted to a hyperbolic tangent function, Eq. 2; *t*_(8)_ = 14, *p* = 7.5 × 10^−7^, two-tailed *t*-test). These patterns are consistent with many previous behavioral results of perceptual tasks (28) and thus suggest that decision-making models similar to those previously proposed, such as bounded evidence accumulation (29), can explain our results.

While performing the task, participants often made multiple saccades between informative features (Fig. 2). Immediately following stimulus onset, their fixations tended to land just below the eyes in the face categorization tasks (Fig. 2A left and middle), which is consistent with previous studies that investigated fixation patterns during face recognition (8). For expression categorization, landing positions appeared slightly closer to the nose (Fig. 2A middle), which also agrees with previous reports (8). For car categorization, the participants’ initial fixation landed on the rear region in most trials. Following this initial fixation, participants often made multiple saccades (Fig. 2D), and their fixation positions were dispersed during decision formation. The density of fixation positions during the full stimulus duration revealed a concentration around the two informative features (Fig. 2B). For identity and car categorization, the density plots exhibited two distinct peaks corresponding to the two features. For expression categorization, the two peaks were less distinct but still covered the two features. These patterns were qualitatively similar across the participants (Supplementary Fig. 3A, B).

**Fig 2.**
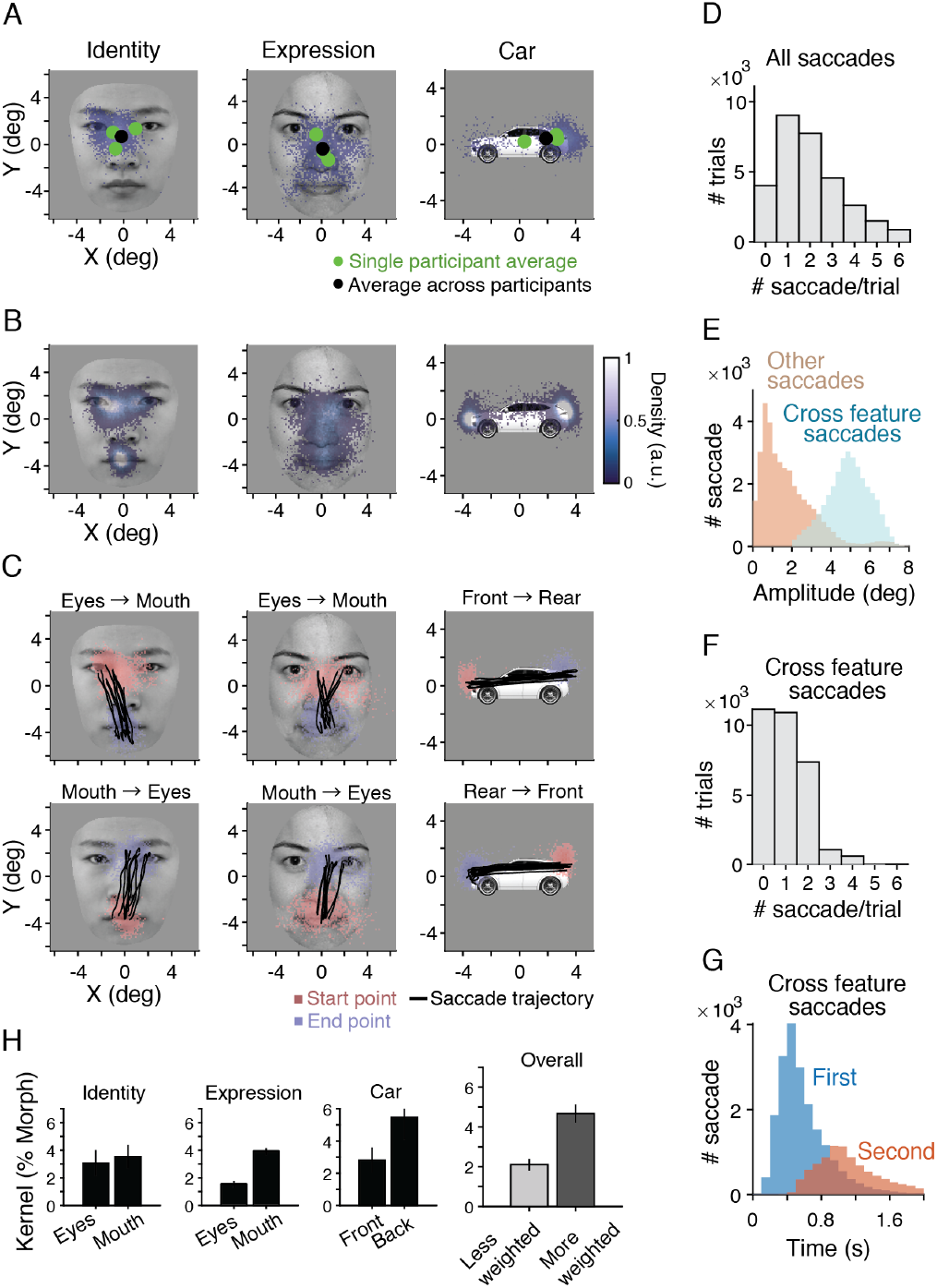
Participants made frequent saccades to sample two informative features. (**A**) Scatter plots of the first fixation positions after stimulus onset show a concentration just below the eyes in the face tasks and the rear part in the car task. The blue dots represent individual trials from a representative participant. Plots for each participant are shown in Supplementary Fig. 3A. (**B**) Density plots of the fixated positions during the entire stimulus viewing period show that the participants primarily fixated around the two informative features. The plots shown are from representative participants; plots for all the participants can be found in Supplementary Fig. 3B. (**C**) Example saccades spanning the two informative features. Plots for each participant are shown in Supplementary Fig. 3C. (**D**) Distribution of saccade counts per trial. The trials were aggregated across participants. Plots for each participant are shown in Supplementary Fig. 4. (**E**) The distribution of saccade amplitudes revealed two peaks. The peak with larger amplitudes corresponded to the saccades spanning the two features (cross-feature saccades; see Methods for its definition). (**F**) Participants made at most two cross-feature saccades in most of the trials. (**G**) Distribution of the timings of cross-feature saccades. (**H**) Consistent with the frequent fixations on the two features, psychophysical reverse correlation (Eq. 6) revealed positive influences of both features on participants’ decisions. Error bars indicate S.E.M. across participants.

**Fig 3.**
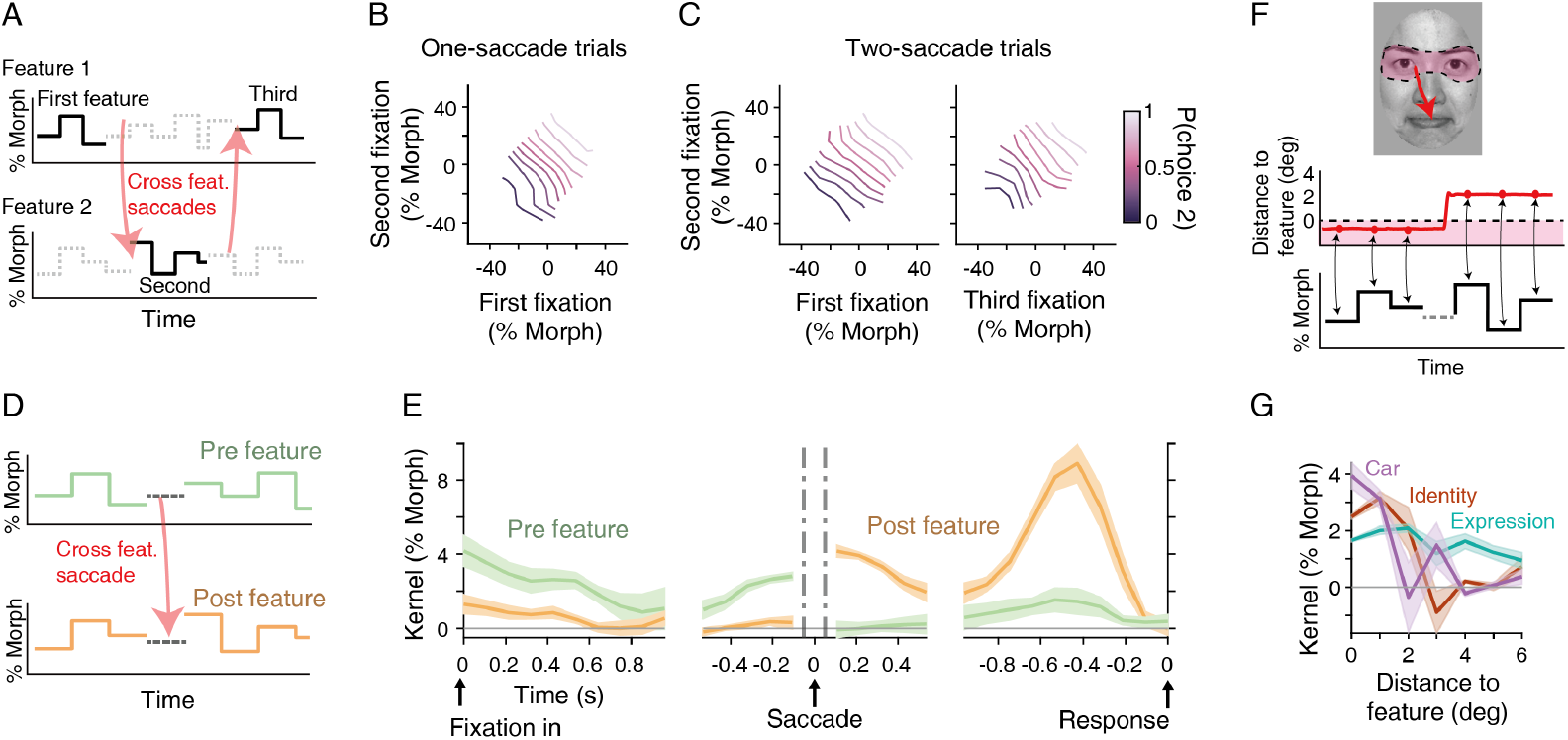
Both features fixated before and after saccades contribute to decisions. (**A**) We extracted the morph levels of the fixated feature (solid lines) at different fixation epochs split by cross-feature saccades and averaged them in each trial for the analysis in B and C. The panel shows an example trial, in which a participant first fixated on feature 1, made a saccade to feature 2, and then fixated back on feature 1. (**B, C**) 2D psychometric functions based on the average morph level of each fixation epoch. The lines are diagonal, indicating that both features fixated across saccades influenced participants’ choices in the trials with one cross-feature saccade (B) and with two cross-feature saccades (C). (**D**) To quantify the temporal weighting of features across saccades, we performed psychophysical reverse correlation (Eq. 6) using the morph fluctuations of the features fixated before and after cross-feature saccades. The schematic shows an example of one-saccade trials. The trials with more saccades are also included in the analysis (see Methods). (**E**) Psychophysical kernels indicate continuous influences of fixated features on participants’ decisions. Their rich temporal dynamics can be explained by a simple evidence accumulation model (see Fig. 4). Shading indicates S.E.M. across participants. (**F**) Schematic for the analysis in G. To examine spatial integration, we sorted the morph fluctuations based on the distance to each feature from participants’ gaze position. The distance was calculated from a contour line manually circumscribing each feature (Supplementary Fig. 1B; see Methods). (**G**) The amplitudes of psychophysical kernels decreased largely monotonically as a function of the distance to the features from the gaze position.

The distinct peaks in the density plots resulted from frequent saccades between the informative features. We plotted the distribution of saccade amplitudes and identified two peaks (Fig. 2E), one corresponding to small saccades within local features and the other to larger saccades spanning across features. We were particularly interested in larger saccades as they lead to large changes in the retinal image and can contribute to the integration of sensory information across distant features. We therefore extracted these “cross-feature” saccades and focused our analysis on their effects on decisionmaking behavior in the following sections. In brief, we defined saccades as cross-features if their start and end points were near the two features and their amplitudes were greater than two degrees (see Methods for more details). Examples of the start and end points of these saccades are shown in Fig. 2C. These cross-feature saccades occurred on average 1.02 ± 0.12 times per trial (Fig. 2F) and appeared to be periodic with an average interval of ∼400 ms (Fig. 2G). We did not consider saccades spanning the left and right eyes as cross-features because the two eyes had the same morph level and did not provide distinct information in our stimuli. The influence of smaller saccades on decision making is shown in Supplementary Fig. 5A.

**Fig 4.**
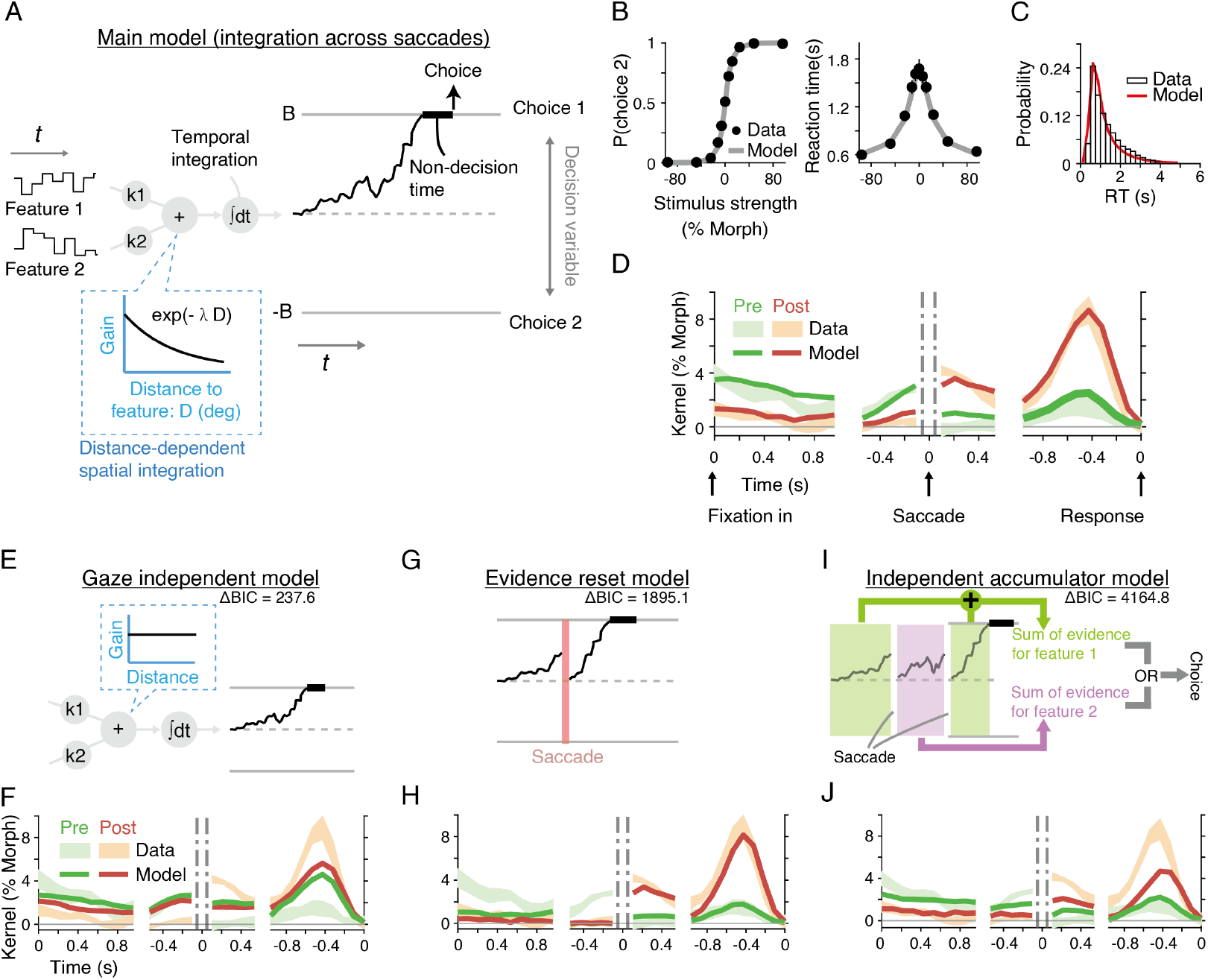
Across-saccade evidence accumulation model explains the behavioral results. (**A**) Our main model accumulates evidence throughout stimulus presentation across saccades until the accumulated evidence (decision variable) reaches a bound. The choice associated with the crossed bound is made after a non-decision time. Momentary evidence is computed as a linear sum of the morph levels of the two features with their weights as free parameters (*k*_1_ and *k*_2_), which are modulated by the distances to the features from the gaze position at each moment (blue inset). (**B, C, D**) The model quantitatively accounts for choices, mean reaction times (RTs), RT distributions (the panel includes the trials of all morph levels), and psychophysical kernels. Plots for individual participants are shown in Supplementary Fig. 6. (**E, F**) If the weights for the features in the model are fixed regardless of the gaze positions (E), the amplitudes of fixated and unfixated kernels become similar, deviating from the data (F). Δ*BIC* indicates the difference in fit performance relative to the main model (positive values indicate poorer fits). (**G, H**) A model that resets evidence accumulation after saccades (G) fails to account for the amplitudes of data kernels before saccades (H). (**I, J**) Alternatively, if a model accumulates evidence independently for each feature and makes a decision based on one of the features (the feature either chosen randomly or based on the magnitude of evidence) (I), it fails to explain the amplitudes of data kernels aligned to the timing of behavioral responses (J).

**Fig 5.**
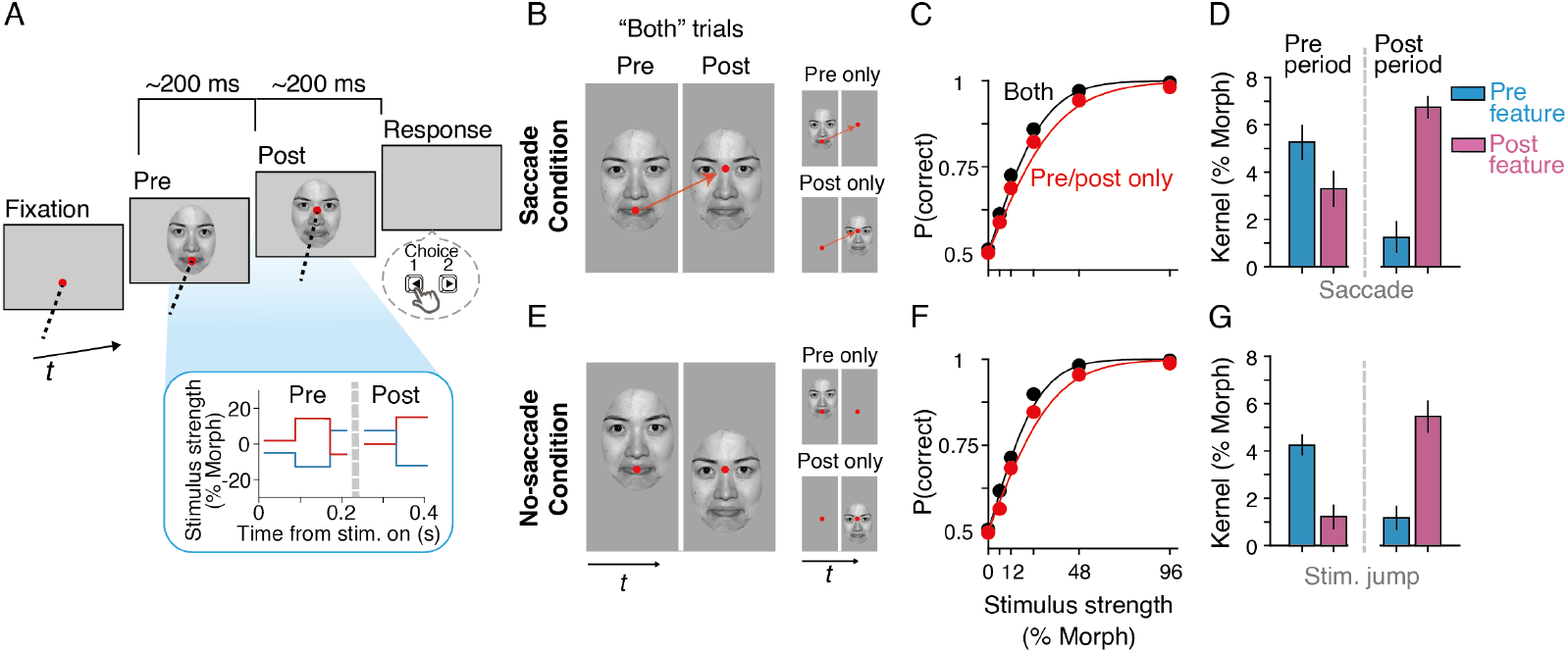
Guided-saccade task revealed a minimum influence of efference copy on feature integration. (**A**) In this task, participants were asked to look at the red fixation dot, which moved from the position of one feature to the other in the saccade condition (B). After the saccade, the stimulus was extinguished in ∼200 ms, and participants reported their choice by pressing a button. As in the free saccade task, morph levels fluctuated every ∼100 ms (inset). Each participant was assigned to perform either identity, expression, or car categorization (*n* = 9). (**B, E**) We compared the saccade condition (B) with the non-saccade condition (E), in which the fixation point stayed at the same position, but the stimulus jumped, mimicking the visual display in the saccade condition. The stimulus duration was matched between the conditions within each participant (see Methods). To quantify the integration of evidence across saccades, we also had trials in which a stimulus was shown only before (“Pre only”) or after (“Post only”) the saccade or stimulus jump event. (**C, F**) Whether or not participants made a saccade, their performance improved when a stimulus was shown in both epochs. Error bars denoting S.E.M. across participants were smaller than the data points. (**D, G**) Psychophysical kernels in the “both” condition showed positive influences of the fixated features before and after a saccade or stimulus jump, consistent with evidence integration. The data of individual participants are shown in Supplementary Fig. 9.

As expected from the frequent saccades between the two features, we observed that participants relied on both of them to judge the stimulus categories. We performed psychophysical reverse correlation (29, 30, 35), in which we averaged the fluctuations of morph levels of each feature during the participants’ viewing of the feature in each trial and computed the difference in the average fluctuations between the trials in which participants chose Category 1 and Category 2 (Eq. 6 in Methods). The amplitudes of the resulting psychophysical kernels quantified the extent to which the fluctuations influenced the participants’ choices (Fig. 2H). The kernels were positive for both features in all the tasks (*t*_(8)_ = 7.1, *p* = 9.9×10^−5^ for the feature weighted less by each participant, *t*_(8)_ = 16, *p* = 2.8×10^−7^ for the feature weighted more, two-tailed paired *t*-test), while the amplitudes often differed between the features such as higher values for mouth than for eyes in the expression task (Fig. 2H). However, such a difference in feature weighting may depend on our choice of prototype stimuli and is not the focus of the current study. Rather, the key observation was that both features were used to solve the task, providing a basis for investigating whether and how eye movements were involved in gathering evidence from distant features.

### Both features fixated before and after saccades contribute to decisions

Having extracted the cross-feature saccades, we now address one of our main questions: whether and how the information of local object features is integrated across these saccades. Many previous studies have demonstrated the trans-saccadic integration of simple visual stimuli (e.g., grating or color patch) seen at the fovea and periphery (14–17, 21, 37, 38), but object recognition poses a different challenge because participants see different features across saccades.

To address this question, we leveraged our stochastic stimuli and first tested whether the morph fluctuations of the features fixated on before and after each saccade correlated with participants’ choices regardless of the features being fixated on. We averaged the morph fluctuations of the fixated features between saccades (Fig. 3A) and plotted the participants’ choice performance as a function of these morph levels (Fig. 3B, C). In the trials with one cross-feature saccade, this became a two-dimensional (2D) psychometric function (Fig. 3B). The plot displayed prominent diagonal iso-performance contours, indicating that both features before and after a saccade influenced the participants’ decisions. Fitting a logistic function (Eq. 4) to this pattern revealed significant weights for both pre-saccade (*t*_(8)_ = 7.15, *p* = 9.7×10^−5^, two-tailed *t*-test across participants) and post-saccade features (*t*_(8)_ = 6.24, *p* = 2.5×10^−4^, two-tailed *t*-test) without a strong interaction term (*t*_(8)_ = −0.39, *p* = 0.70,BF_01_ = 2.91, two-tailed *t*-test). Likewise, we analyzed the trials with two cross-feature saccades and confirmed that the morph levels of the features fixated during the first, second, and third fixation periods all correlated with the participants’ choices (Fig. 3C; first period: *t*_(8)_ = 3.66, *p* = 0.0064, second period: *t*_(8)_ = 9.13, *p* = 1.7×10^−5^, third period: *t*_(8)_ = 4.57, *p* = 0.0018, interactions: *p* > 0.09,BF_01_ = 2.64, two-tailed *t*-test; Eq. 5). Thus, the participants relied on information both before and after saccades to make their decisions. In the next section, we show that these results indicate the integration of evidence rather than random reliance on features before or after saccades.

We then used psychophysical reverse correlation to quantify the temporal dynamics of feature weighting and found a persistent contribution of fixated features across saccades. We computed the psychophysical kernels over time by calculating the difference in stimulus fluctuations at each time point between the trials in which the participants chose Category 1 and Category 2 (Eq. 6). The resulting kernels revealed rich temporal dynamics (Fig. 3E) and, importantly, had positive weights throughout the stimulus presentation when the feature was fixated (Fig. 3E; “pre” and “post” features indicate the features fixated before and after a saccade, see Fig. 3D). When aligned to stimulus onset, the kernels tended to gradually decrease over time. Around the time of cross-feature saccades, the amplitudes of the pre- and post-saccade features were swapped. Because the temporal resolution of our stimulus fluctuations was ∼100 ms, we did not analyze further details of temporal dynamics around saccades (cf. (15); but see Supplementary Fig. 5B for kernels plotted with a higher temporal resolution). When aligned to the time of the participants’ choice, we observed a charac-teristic peak abound 400-500 ms before the choice (Fig. 3E right). As demonstrated in the next section, these complex kernel dynamics can be explained quantitatively using a simple evidence accumulation model.

Psychophysical reverse correlations could also be used to quantify the spatial integration of sensory evidence. Instead of sorting stimulus fluctuations over time, we sorted the same data according to the visual distance between the participant’s gaze position and each feature at each time point (Fig. 3F; see Methods for the definition of distance). The resulting kernels revealed a largely monotonic reduction as a function of distance (Fig. 3G) with some variability across the stimulus conditions. In the identity and car tasks, kernel amplitudes decreased sharply with distance, whereas they were much flatter in the expression task. Consistent with this finding, another line of analysis (Supplementary Fig. 8A, B) confirmed that the influence of unfixated features was markedly greater on the expression task. Thus, the extent of the spatial window for integration can be either stimulus-or task-dependent (see Discussion). Nonetheless, the influence of the features can still be described as a monotonic function of the visual distance from the features in all tasks.

### Across-saccade evidence accumulation accounts for behavior

Encouraged by our observation of the positive influence of features fixated on before and after saccades, we formally tested the integration of evidence across saccades by fitting an evidence accumulation model to the behavioral data. Previous studies have shown that face categorization behavior during fixation conditions could be explained using a simple model that computes the weighted sum of evidence from each facial feature and accumulates this sum over time (29, 30). We extended this model by incorporating participants’ eye movements such that the model kept accumulating evidence across saccades, but the informativeness of each feature depended on the gaze position (39).

Our model integrates fluctuating sensory evidence from object features (e.g., eyes and mouth in face tasks) and accumulates evidence over time across saccades to form a decision variable (Fig. 4A). Each feature has a different strength of evidence (sensitivity parameter *k*_*i*_ in Eq. 10), which de-cays as a function of the distance between the feature and the participant’s gaze position at each time point (decay rate λ in Eq. 10). Decay was modeled using an exponential function based on previous studies (8, 40), but other monotonic functions could similarly fit the data (Supplementary Fig. 7B). Aside from this decay, we did not assume any component in the model that depended on eye positions and saccades. When the decision variable reaches an upper or lower bound, the model makes a choice associated with the bound after a non-decision time that accounts for sensory and motor delays.

This simple extension of an evidence accumulation model accounted for all aspects of the behavioral data examined. The model accurately fitted the choices, mean RTs, and the distributions of RTs (Fig. 4B, C; *R*^2^ = 0.83 ± 0.028). Furthermore, it quantitatively accounted for the patterns of psychophysical kernels aligned to all of the time epochs (Fig. 4D; *R*^2^ = 0.51 ± 0.074). Note that the model kernels were not directly fitted to the participants’ kernels but were simulated from an independent set of stimulus fluctuations (see Methods), making the comparison of data and models informative. The dynamics of kernels observed in the data can be accounted for by the mechanistic components of evidence accumulation (29, 30). The model explains the decreasing kernels aligned to stimulus onset because there is a temporal gap between the bound crossing and the report of a decision (i.e., the non-decision time), making a later portion of the stimulus fluctuations irrelevant to the decision. Because the timing of the bound crossing varies across trials, the model predicts a gradual reduction in the effect of stimulus fluctuations over time. The peak of the kernels aligned to the behavioral responses corresponded to the moment of crossing a decision bound in the model. At that moment, even tiny stimulus fluctuations bring the decision variable beyond a bound and dictate the decision, leading to the large kernel amplitudes. Subsequently, the kernels sharply drop to zero because of the non-decision time. When aligned to the time of saccades, the amplitudes of the kernels swapped between the pre- and post-saccade features because of the change in sensory sensitivity caused by the distance between the features and the gaze location.

We further confirmed that no other model accounted for the behavioral data without assuming gaze-dependent ev-idence accumulation. First, we tested a model that did not consider gaze position but accumulated evidence from two informative features with constant sensory sensitivity over time (Fig. 4E). This gaze-independent model could fit choices and RTs (Supplementary Fig. 7C) but clearly failed to explain the differences in the amplitudes of psychophysical kernels between fixated and unfixated features (Fig. 4F). The model kernels showed slight differences between fixated and unfixated features because participants tended to fixate on features with higher sensitivity more frequently, but the differences were far smaller than those observed in the actual data.

Second, we considered a model that did not integrate evidence across saccades. One plausible scenario is that participants restarted evidence accumulation after each saccade (Fig. 4G); if a decision could not be made based on one feature, they switched their focus to the other feature and made a decision based on it. This model could also fit choices and RTs (Supplementary Fig. 7D), but, as expected, failed to explain the amplitude of psychophysical kernels before saccades (Fig. 4H). The model kernels before saccades were slightly positive owing to the inclusion of trials without saccades, but they were much smaller than the data.

Finally, we ruled out a hypothesis that participants relied on either feature before or after saccades to render a decision but did not integrate evidence across saccades (Fig. 4I). This independent accumulator model computed the total sum of evidence for each feature while viewing that feature and then committed to a choice using the evidence from one of the features at the moment of the decision. One variant of this model chose a feature randomly, with the probability of choosing the last fixed feature as a free parameter (Fig. 4J, Supplementary Fig. 7E). Another variant chose the feature with a higher magnitude of evidence at the moment of the decision (Supplementary Fig. 7F). Because both features before and after saccades could contribute to decisions in different trials, the kernels of these models had positive amplitudes throughout the stimulus presentation (Fig. 4J). However, the model kernels had a smaller peak aligned to the participants’ responses because the bound crossed during the last fixed feature did not always dictate the decisions. This contrasts with our main model, in which the crossed bound always corresponded to the model’s decisions. Our data clearly favor the main model.

Overall, we found that a simple mechanism that accumulates sensory evidence across saccades is sufficient to account for the participants’ object categorization behavior. Eye position information was required in the model to explain the decrease in sensitivity to features as a function of visual eccentricity; however, aside from that, we did not need to model the complex interactions between eye movements and the object-recognition process. While these results cannot prove the absence of such interactions, our results in the next section also support the idea that the interactions of the visual and oculomotor systems do not play a substantial role in the types of object recognition behavior we examined.

### Active saccade commands are unnecessary for feature integration

The observed integration of evidence across saccades could have depended on neural processes that combine visual signals and active saccade commands (i.e., efference copy), such as the predictive coding of visual features prior to saccades. To examine the extent of the contribution of efference copy, we next designed a “guided-saccade” task that could compare object recognition performances between conditions with and without saccades (Fig. 5A). This task used the same object stimuli as in the free saccade task, but we strictly controlled the participants’ eye movement and stimulus presentation. In the saccade condition, participants were explicitly instructed to make cross-feature saccades during object categorization (Fig. 5B). In the no-saccade condition, participants maintained fixation while there was a sudden change in the visual display, mimicking the change caused by saccades (Fig. 5E).

The saccade and no-saccade conditions had almost identical trial structures and stimulus durations. In each trial, a fixation point was initially placed at the center of one of the informative features of the stimulus. In the saccade condition, the fixation point moved to the location of the other informative feature immediately following the stimulus onset (Fig. 5B). The participants were required to make a saccade following this jump of the fixation point. The saccade was followed by another stimulus period (∼200 ms), and the participants reported their decisions by pressing a button after the stimulus was extinguished. In the no-saccade condition, the stimulus position suddenly shifted from one region to another, while the fixation point stayed the same, and the participants had to maintain fixation (Fig. 5E). The duration of the first stimulus and the blank between the two displays were set according to each participant’s saccade latency (170.0 ms ±5.3 ms) and duration (53.5 ms ±1.3ms) in the saccade condition; thus both the spatial and temporal profiles of stimuli were approximately matched between the two conditions. Each of the two conditions had trials where a stimulus was shown both before and after a saccade/stimulus jump (“Both” trials; Fig. 5B, E left) and trials where a stimulus was shown only before or after (“Pre only” and “Post only” trials; Fig. 5B, E right).

In the saccade condition, we confirmed that participants integrated evidence across a saccade. Their behavioral accuracy was significantly higher when an image was present across a saccade (“Both” trials) than when it was present only before or after a saccade (Fig. 5C; the difference in logistic slope *α*_2_ = 1.69 ± 0.36, Eq. 3; *t*_(8)_ = 4.75, *p* = 0.0014, two-tailed *t*-test). The improvement in performance was subtle but consistent with the near-optimal integration of evidence (Supplementary Fig. 9), in line with prior studies that examined the saccadic integration of simpler features (14). Similar to the free-saccade task, we also identified positive kernels for the fixated feature both before and after a saccade (Fig. 5D; before saccade *t*_(8)_ = 6.41, *p* = 2.1 × 10^−4^, after saccade *t*_(8)_ = 10.8, *p* = 4.8 × 10^−6^, two-tailed *t*-test).

Critically, higher performance for the “Both” trials was also observed in the no-saccade condition, supporting evidence integration (Fig. 5F; the difference in logistic slope *α*_2_ = 2.01±0.39, Eq. 3; *t*_(8)_ = 5.12, *p* = 9.1×10^−4^, two-tailed *t*-test). In support of this, we also identified the positive kernels for the fixated features both before and after a saccade in the no-saccade condition (Fig. 5G; before saccade *t*_(8)_ = 9.69, *p* = 1.1×10^−5^, after saccade *t*_(8)_ = 7.97, *p* = 4.5×10^−5^, two-tailed *t*-test). The size of the performance improvement was statistically indistinguishable from the saccade condition (*α*_2_ = 0.32±0.50, Eq. 3; *t*_(8)_ = −0.64, *p* = 0.54, BF_10_ = 0.38, two-tailed *t*-test). One potential difference we noted was that the kernel for the unfixated feature before a saccade looked slightly higher in the saccade condition than in the no-saccade condition, which may indicate pre-saccadic enhancement in visual sensitivity (41), but this difference was also statistically indistinguishable (Fig. 5D, G; *t*_(8)_ = −1.71, *p* = 0.13, BF_10_ = 0.93, two-tailed *t*-test). Thus, the results indicate that saccade commands are not a prerequisite for feature integration and do not substantially improve behavioral performance even if they are effective (see Discussion for the interpretation).

### Saccade frequency was minimally influenced by ongoing decision formation

Thus far, we have focused on the mechanisms of perceptual decision making during object recognition, but our free saccade task (Fig. 1) also allowed us to examine whether and how saccade patterns are modulated by the ongoing decision-making process. For example, when the currently fixated feature is uninformative, people may make frequent saccades for the other feature to seek for more evidence (42), or people may spend more time fixating on each feature (43, 44). If so, there could be a correlation between stimulus difficulty and saccadic frequency.

Contrary to these expectations, we did not find any significant relationship between stimulus difficulty and saccade frequency in our tasks. The average number of cross-feature and other saccades in each trial was clearly higher in more difficult trials (Fig. 6A), but this did not indicate more frequent saccades because the trial duration (i.e., reaction time) was longer in difficult trials (Fig. 1C bottom). Therefore, the number of saccades per time had to be estimated, but the calculation requires greater complexity than a simple division of the number of saccades by the trial duration, as saccades tend to be periodic (Fig. 2G), and the calculation strongly depends on the relative distributions of RTs and saccade timing. Therefore, we matched the RT histograms of different stimulus strengths by randomly subsampling trials. After the RT matching, saccade frequency was not correlated with stimulus difficulty (Fig. 6B; cross-feature saccades: *F*_(5,48)_ = 0.08, *p* = 0.995, BF_01_ = 9.26; all saccades: *F*_(5,48)_ = 0.1, *p* = 0.992, BF_01_ = 7.52, repeated-measures one-way ANOVA).

**Fig 6.**
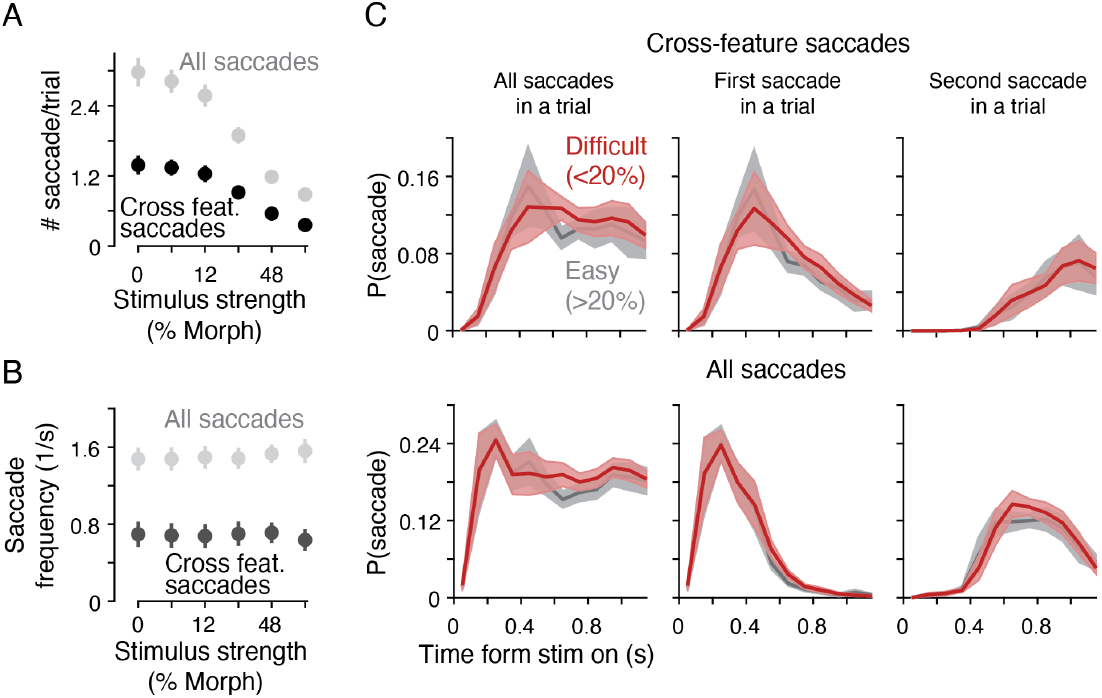
The frequency of saccades did not significantly depend on stimulus difficulty. (**A**) The average number of saccades per trial was higher for more difficult (low morph level) trials. Error bars indicate the S.E.M. across participants. But reaction times were also longer for these trials (see Fig. 1C). (**B**) When saccade frequency per time was calculated, it was not correlated with stimulus difficulty. To allow the unbiased estimation of saccade frequency, we matched RT distributions across stimulus strength when calculating the frequency (see main text and Methods). Plots for each participant are shown in Supplementary Fig. 10A. (**C**) Without matching RT distributions, we compared the probability of making a saccade at each time point by dividing the count of trials with a saccade at that time point by the count of trials whose RTs were longer than that time. We compared this probability between easy and difficult trials and found that they were statistically indistinguishable. Plots for each participant are shown in Supplementary Fig. 10B.

We further corroborated this conclusion through an analysis that does not rely on matching RTs (Fig. 6C). This was important as RTs could be highly correlated with participants’ subjective uncertainty (45) and thus matching RTs could also align subjective uncertainty across stimulus strengths. Instead, we calculated the probability of making a saccade at each time point using only trials with RTs longer than the time point. As long as there were a sufficient number of trials at each time point, this probability should not be biased by the duration of the RTs following each time point. We plotted this probability using either all cross-feature saccades, the first cross-feature saccade in a trial, or the second saccade in a trial, and did not find differences between easy and difficult trials (Fig. 6C top; split by 20% morph boundary; *p* > 0.7, BF_01_ > 10, repeated-measures two-way ANOVA). Including within-feature saccades did not affect this result (Fig. 6C bottom; *p* > 0.5, BF_01_ > 10, repeated-measures two-way ANOVA). In summary, saccades appeared to occur in a stochastic manner without clear influence from the ongoing decision-making process in our tasks.

## Discussion

Humans often make saccades to sample local informative features when viewing object images, but the mechanisms by which saccades contribute to object recognition have yet to be established. Studies on eye movements have proposed complex interactions between the visual and oculomotor systems, such as predictive visual processing based on saccadic commands (26), whereas studies on object recognition lean toward eschewing this complexity and favor a briefly flashed static image under fixation conditions (46). Here, we applied a decision-making theory to reformulate the problem as the accumulation of sensory evidence from multiple local features across saccades. Our results indicate a parsimonious relationship between eye movements and object recognition; humans integrated evidence across saccades (Figs. 3, 4), but behavioral performance did not strongly depend on active saccade signals (Fig. 5). As such, a simple evidence accumulation model that does not assume complex interactions between the visual and oculomotor systems can approximate decision-making behaviors.

Many prior studies have documented the integration of visual information across saccades (14–17, 21, 37, 38), but to our knowledge, ours is the first attempt to apply a mechanistic model grounded on evidence accumulation to account for object recognition involving saccades. The accumulation of evidence is common in many perceptual tasks (28), and existing models of form and object vision suggest that information is integrated across saccades (23–25). Thus, one might argue that our results were largely expected. However, we believe that we have provided important empirical tests for the following three points. First, we were able to demonstrate the integration of evidence across saccades using naturalistic stimuli (faces and objects) through a modelfitting approach that quantitatively explains various aspects of the participants’ choice behavior. Second, this modeling approach allowed us to test the integration process during a free viewing condition in which participants’ gaze locations and times were unrestricted. Finally, we demonstrated a limited role for efference copy in this process, confirming that a simple evidence accumulation model (Fig. 4) is sufficient to explain behavior in our tasks.

The apparent lack of necessity for efference copy (Fig. 5) seemingly contradicts a large body of existing studies underscoring its role in visual perception (26, 27, 47, 48), but this could stem from the fact that the demands for our tasks were different. In conventional trans-saccadic perceptual tasks, participants are required to judge a simple stimulus (e.g., the orientation of a Gabor patch) initially viewed in the periphery and subsequently foveated through a saccade (14–17). Here, the remapping of receptive fields induced by oculomotor commands (26) could serve as a vehicle to fuse information across saccades (27, 47), although the necessity of an efference copy is still debated even under this condition (49, 50). In contrast, object images comprise multiple distinct features. We believe that the integration of foveated features across saccades is more important for improving behavioral accuracy in this case, which may not require oculomotor information. It is also possible that our behavioral data did not have sufficient statistical power to detect the effects of oculomotor signals on feature integration, but even if there is an effect, its effect size must be limited according to our results.

Regarding the frequency of saccades, we found minimal influence of uncertainty in ongoing decision making (Fig. 6). Previous studies have found that the timing of saccades is modulated by internal states of visual recognition (44, 51–56). We did not observe such effects in our tasks, possibly because our participants were extensively trained to perform saccadic sampling on the same visual images and had established stereotyped saccadic patterns based on the learned statistics of the stimuli. Also, the average stimulus strength was always matched between two informative features in our tasks, potentially providing less incentive for participants to modulate their saccade patterns depending on decision uncertainty. Nevertheless, the lack of effects suggests that evidence accumulation during decision making and eye movement planning can be independent. Some previous theories have used the drift-diffusion process to explain saccade timing (51, 55, 57), but it is likely that such a process and the evidence accumulation we modeled here are separate processes in the brain.

The spatial distribution of saccades we observed were consistent with previous studies (Fig. 2). Participants tended to look at informative regions or nearby areas in all tasks (Fig. 2B) and made frequent saccades across features (Fig. 2C). A seminal study by Peterson and Eckstein (2012) showed that people look just below the eyes during face recognition, which is consistent with the prediction of an ideal-observer model. Our observations of the initial fixation positions replicated these findings (Fig. 2A), whereas fixations diverged to other locations during prolonged viewing, which is also consistent with their results (58). A tantalizing question is whether this behavior is optimal in terms of information sampling (59–61), but defining optimality in our tasks is not trivial. An ideal-observer model may recommend maintaining fixation on the most informative part of an image, but sampling multiple informative features might also be optimal depending on its definition (24, 25, 62).

It would also be worthwhile to discuss how our findings extend to other natural object stimuli. We designed our stimuli to have two distant informative features to manipulate informativeness and detect the participants’ saccades across the features. Although we designed our stimuli to be naturalistic—eyes and mouth are indeed informative features for face recognition (29, 63)—other object images would have more than two diagnostic features that could also overlap with each other. In such cases, people may not directly look at each feature but rather look at a place between the features to sample evidence. Indeed, in the expression task, the participants tended to fixate between the eyes and mouth (Fig. 2B) and had a broader spatial window (Fig. 3G) sampling evidence from both fixated and unfixated features (Supplementary Fig. 8). This might indicate that the participants adjusted their spatial sampling windows depending on the task context. Stimulus size would also influence sampling strategies (1, 64). In our experiments, we used a naturalistic range of object sizes (36), but if the viewing distance increases, the image would become too small to make saccades inside, in which case features could be spatially integrated without saccades (29). Although we were unable to test these diverse conditions, our model —evidence accumulation that depends only on the eccentricity of features— can readily offer quantitative predictions of object-recognition behaviors in any of these settings.

We hope that the simplicity of the proposed framework encourages further investigation of object recognition under naturalistic viewing. Our findings suggest that, despite the apparent complexity of oculomotor events, there could be stable representations of momentary and accumulated sensory evidence across saccades in the neural circuitry for object processing (65, 66) and decision making (67). Such a solution for active object vision can also be easily implemented in image-computable models, since it only requires accumulating evidence from the model responses to each snapshot of image sequences.

## Methods

### Participants and experimental setup

We recruited 22 participants (age 20-40, 4 males and 18 females, students or employees of the Chinese Academy of Sciences). Our participant sampling did not consider sex, as it was deemed unlikely to influence the outcomes of this study. All participants had normal or corrected-to-normal vision and were naive to the purpose of the experiment. Written informed consent was obtained from all participants. Six of them were dropped before the main data collection, either because of scheduling issues or poor eye-tracking quality. All experimental procedures were approved by the Institutional Review Board of the Center for Excellence in Brain Science and Intelligence Technology (Institute of Neuroscience), Chinese Academy of Sciences.

Nine participants performed the free-saccade task (Fig. 1), while nine performed the guided-saccade task (Fig. 5). Two of them performed both tasks. In each task, we assigned participants to perform categorization of face identity stimuli, face expression stimuli, or car stimuli (Fig. 1A; three participants each), but this study focused on the results consistently observed across these stimulus conditions. Sample sizes were chosen following the convention of studies using psychophysical reverse correlations and modeling of decision-making behavior (29). We had to set our sample sizes small because we sought to collect a large number of trials from each participant (∼3,500 trials per participant; 63,984 total trials in this study) after extensive practice sessions (∼2,000 training trials per participant prior to data collection) in order to obtain as much reliable behavioral data as possible from individual participants (68). We performed a sensitivity analysis and estimated that our sample size (9) could detect an effect with 80% power provided the standard deviation across participants was less than 90% of the effect size (69). When the statistical tests did not show significance, we supplemented our analysis with the Bayes factor and interpreted the results cautiously.

During the experiments, the participants sat in a height-adjustable chair in a semi-dark room. Their chins and foreheads were supported by a chin-rest mounted on a table, which was fixed at a specific position to ensure a stable viewing distance of 57 cm from a cathode-ray-tube monitor (17-inch IBM P77; 75 Hz refresh rate; 1024 × 768 pixel screen resolution). The Psychophysics Toolbox (70) and MATLAB (MathWorks, Natick, MA, USA) were used to control the stimulus presentation. The eye movements were monitored using a high-speed infrared camera (Eyelink 1000 Plus; SR Research, Ottawa, Canada). The gaze position was recorded at 1 kHz.

### Task designs

#### Face and object categorization task with free eye movement

To examine the contribution of saccades to object recognition, we designed three versions of the object categorization task: face identity, face expression, and car categorization (Fig. 1A). In each version of the task, participants classified an image into one of two categories while they were allowed to freely make saccades inside the image. The stimuli were chosen from a morph continuum created by interpolating two prototype images, and the participants were required to report which prototype the given stimulus looked similar to. The two prototypes were male and female faces in the identity task, happy and sad faces of the same individual in the expression task, and two types of cars in the car task (Fig. 1A). We used face stimuli because they allow easy definition of informative features (i.e., eyes and mouth) and previous studies have successfully explained face categorization during fixation conditions using an evidence accumulation model (29, 30). We further designed a car categorization task to ensure that the results were not specific to face recognition.

Participants began each trial by fixating on a fixation point (0.3° diameter), which appeared randomly at one of six peripheral locations (11.5° away from the screen center for the face tasks, 8° for the car task) (Fig. 1B). After a short delay (400-700 ms, truncated exponential distribution), a stimulus appeared at the center of the screen. The randomized fixation point locations were intended to minimize bias in the location that participants initially looked at in the image (8, 71). Participants then had to make a saccade in the stimulus within 500 ms of its appearance. The stimulus was kept ambiguous (halfway between the two prototypes on the morph continuum) and thus uninformative until the participants made a saccade. After fixation, the stimulus was replaced with a face image of the morph level chosen for the trial. The participants were then allowed to look at any place in the image, but if their fixation left the image, the trial was aborted. Participants reported the category of the stimulus by pressing one of two keyboard buttons whenever they were ready (reaction-time task). The stimulus was extinguished immediately when the button was pressed. If the participants did not make a decision within 5 s, the trial was aborted. In total, 1.75% of trials were aborted either due to fixation breaks during stimulus presentation or time out. Auditory feedback was provided for correct and incorrect decisions. If the stimulus was ambiguous (halfway between the two prototypes on the morph continuum), the correct feedback was provided in a random half of the trials. Following feedback, the next trial began after a 1 s inter-trial interval.

We created each stimulus set by continuously morphing two prototype images using a custom-made program (29). Prototypes for the expression task were obtained from the Nim Face set (33), prototypes for the identity task were obtained from the Tsinghua Facial Expression Database (34). The face images shown in the figures were from these databases and presented with permission. The prototypes for the car task were generated by authors using Midjourney (https://www.midjourney.com) using prompts such as “Clean and minimalist product photography of a white SUV with soft edges, highlighting its sleek and modern design”. We then used Photoshop (Adobe, San Jose, CA) to edit the image parts that were difficult to morph, such as the steering wheels, to create a naturalistic morph continuum. Our program generated the morphed intermediates of the two prototypes by linearly interpolating the positions of manually defined anchor points on the prototypes and textures within the tessellated triangles defined by the anchor points. The linear weights for the two prototypes determined the morph level of an image (ranging from -100 to 100%, where the two extremes corresponded to the two prototypes). In each trial, we chose an average morph level from -96, -48, -24, -12, -6, 0, 6, 12, 24, 48, 96%. For participants with higher performance, we also added -3 and 3% morph levels.

Our algorithm could morph local stimulus features independently. For face images, we manually circumscribed the regions containing the eyes and mouth and morphed only the inside of the regions (Fig. 1A). Similarly, for the car images, we manually defined the front and rear regions (Fig. 1A). The regions outside these features were maintained halfway between the two prototypes and thus remained uninformative. For face images, because the regions outside the eyes and mouth show a limited contribution to judgments (29, 63), this segmentation of informative and uninformative regions was unlikely to influence participants’ behavior. For the car images, the front and rear parts were split at the midline (Fig. 1A), but the two prototypes were mostly different around the hood (bonnet) and the rear window. While the choice of prototype images would affect which parts become informative, we do not consider that this choice affected the main conclusions of this study. We could also independently adjust the full dynamic range of the morph line for each feature. This adjustment was made when we realized that the participant relied heavily on (kept looking at) one feature during training. We gradually decreased the range of the over-sampled features up to 50% while confirming that the participant’s overall performance was maintained. Once the main data collection began, we did not make any adjustments.

To examine how each object feature contributed to the participants’ decisions, we added random temporal fluctuations of the morph level to the individual features in each trial. The mean morph level was fixed within a trial and matched between the two features, but the morph level of a feature was randomly updated every 106.7 ms (eight monitor frames of the 75Hz display) drawn from a Gaussian distribution with an SD of 20%. The 106.7 ms fluctuation duration provided us with sufficiently precise measurements of the participants’ temporal weighting in their ∼1 s decision time, while the duration was sufficiently long to ensure a subliminal transition of the morph levels from one image to another. Between two morphed face images, we interleaved a noise mask (phase randomization of the 0% morph face) with a smooth, half-cosine transition function during the eight monitor frames (29). This mask minimized the chance that participants noticed fluctuations in morph levels during stimulus presentation. An example movie of the dynamic face stimuli can be found in Okazawa et al. (2021).

In each trial, the stimulus fluctuations started only after the participants made a saccade into the stimulus. Prior to the saccade, participants fixated on a fixation point placed peripherally (8-11.5° away from the image). During this period, the stimulus remained uninformative (0% morph). Thus, the participants could start judging the stimulus category only after fixating on it. Exactly at the moment of saccade to a stimulus, the stimulus underwent a sudden change in the morph level, but no participants noticed this change. Since the stimulus fluctuations started only after fixation on the image, psychophysical kernels (Figs. 3E, 4) were aligned to the timing of the participant’s fixation on the stimulus rather than the timing of the actual stimulus onset.

We determined the stimulus sizes in our tasks to characterize human object recognition behavior under natural conditions. We thus set the distance between the two informative features (eyes and mouth for face stimuli, and front and rear parts of car stimuli) to be five visual degrees apart. This is approximately the size that we experience when seeing objects and faces at natural distances (36). Under this constraint, the full stimulus size was ∼9.3° ×11° (W H) for the identity task, ∼9.4° ×13.6° for the expression task, and ∼7° ×2.8° for the car task. We expect that participants’ saccade patterns would be greatly affected by stimulus size (1, 64). For example, participants would cease making saccades when the stimulus becomes too small. However, our primary goal was to study the effect of saccades under naturalistic conditions, where saccades would spontaneously occur, and we believe that our conclusions hold as long as they are tested under such naturalistic ranges (see Discussion).

We recruited nine participants for the free saccade tasks, of whom three each were randomly assigned to perform each of the three categorization tasks (31,128 trials in total; 3,459 ± 106 trials per participant). Prior to the main data collection, the participants underwent extensive training (on average 2,000 trials) to ensure stable behavioral accuracy. During training, we informed the participants that the images contained multiple informative features for categorization and encouraged them to use multiple features to solve the task. However, we did not directly ask them to make eye movements or to look at particular parts of an image.

#### Guided saccade task

To examine whether oculomotor commands are necessary for feature integration, we designed a guided saccade task in which participants categorized objects with or without a saccade (Fig. 5A). In the saccade condition (Fig. 5B), we instructed participants to make a saccade during the stimulus presentation by moving the fixation point from one region to another. In the no-saccade condition (Fig. 5E), participants maintained fixation while the stimulus position suddenly moved, mimicking the change in retinal input resulting from a saccade. Similar to the free saccade task, we used face identity, expression, and car categorization conditions with the same stimuli and categorization rules.

In the saccade condition, participants initially viewed a fixation point that appeared at one of two locations near the center of the screen. The two locations corresponded to the location of the two informative features of a stimulus shown shortly afterward (eyes or mouth in the face tasks, front or rear in the car task). There was a variable delay between the participant’s fixation onset and stimulus onset (400-700 ms, truncated exponential distribution). Immediately after the stimulus onset, the fixation point shifted to the location of the other informative feature, and participants had to make a saccade to this location in between 100-400 ms. After sufficient training, participants could consistently make a saccade with ∼200 ms latency (timeout: 5.1% of trials). After the saccade, the stimulus continued for another 213.4 ms; thereafter, it disappeared together with the fixation point, and the participants had to report their decision by pressing a keyboard button within 1 s (timeout: 1.5% of trials). These brief stimulus presentations replicated previous studies that investigated trans-saccadic integration (14) and were ideal for testing the temporal integration of evidence because behavioral performance is likely to saturate with longer stimulus duration (72).

In separate blocks, we performed the no-saccade condition, which mimicked the change in retinal input during the saccade condition without asking participants to make an actual saccade (Fig. 5E). In this condition, the fixation point remained in the same place, but a stimulus briefly appeared with one feature centered at the fixation point, and after a brief blank period, it reappeared with the other feature centered at the fixation point. Thus, the condition approximated what the participants would have seen during the saccade condition. The duration of the first stimulus presentation and the blank were randomly sampled from the distribution of the saccade latency (170.0 ms ± 5.3 ms) and saccade duration (53.5 ms ±1.3ms) obtained in the main condition for each participant. To obtain these numbers, we first collected half of the data for the saccade condition. Subsequently, we collected the remaining half together with the no-saccade condition in the same sessions to ensure that each participant’s training level was similar between the two conditions.

In the saccade and no-saccade blocks, we also included trials in which a stimulus was shown only before or after a saccade/stimulus jump (Fig. 5B, E). In these trials, we removed/displayed a stimulus contingent on the timing of the participant’s saccade (either “Pre only” or “Post only” trials in Fig. 5B, E right). These trials were randomly interleaved with the main condition in which a stimulus was shown in both periods (“Both” trials; Fig. 5B, E left). This allowed us to test whether participants’ performance improved when a stimulus was present both before and after a saccade, indicating the integration of evidence across saccades.

The stimulus morph levels fluctuated in this task, similar to those in the free saccade task. Because the morph levels were updated every 106.7 ms (eight monitor frames, during which a stimulus image made a smooth transition to a noise mask; see above), approximately two cycles of fluctuations occurred before a saccade, as the saccade latency was, on average, ∼170 ms. After the saccade, we reset the fluctuation cycle such that one cycle started immediately after the saccade landing, ensuring consistency in the pattern of stimulus-mask cycles before and after the saccade. The post-saccade stimulus fluctuations continued for two cycles (213.4 ms) before the stimulus was terminated. As in the free saccade task, fluctuations occurred independently for the two informative features, whereas the average morph level was the same for the two features and was constant during the trial. The average morph levels were chosen from -96, -48, -24, -12, -6, 0, 6, 12, 24, 48, 96%. For participants with higher performance, we added -3 and 3% morph levels.

Nine participants performed this guided saccade task, of whom three each was randomly assigned to the facial identity, expression, or car categorization conditions (32,856 trials in total; 3,651± 68 trials per participant). Prior to the main data collection, the participants underwent extensive training (on average 2,000 trials) to ensure stable saccade latency and behavioral accuracy.

### Data analysis

#### Detection of cross-feature saccades and quantification of saccade patterns

Saccades were detected from the 1 kHz eye tracking data using the Eyelink 1000 Plus’ default saccade detection parameters with additional criteria (73) to ensure accuracy. We first applied a small smoothing (a Gaussian filter with 3 ms standard deviation) to the tracking data to remove high-frequency noise and then detected the timings of eye traces with a velocity exceeding 30 °*/s* for 4ms and acceleration exceeding 8000 °*/s*^2^ for 2ms. These timings were considered potential saccade onsets. We then estimated the end time of these potential saccades by looking for the time when the velocity fell below 20 °*/s* for 2 ms. Finally, we classified them as saccades if their duration was longer than 6 ms and they were observed at least 20 ms after the last saccade (73). Through manual inspection, we confirmed that these parameters accurately detected saccades. In rare cases, noise in the eye traces led to the false detection of saccades, which were removed during manual inspection.

Our key objective was to examine how large saccades spanning multiple features in an image contribute to the integration of evidence across features. We thus focused on these cross-feature saccades in our main analyses. We considered a saccade a cross-feature if it satisfied the following criteria. (1) The saccade start point was inside or near the region of one feature (< 1.5° for the face tasks and < 0.5° for the car task) and its end point was inside or near the region of the other feature. These numbers were chosen based on manual inspection of eye movement patterns. The region for each feature was manually circumscribed (Supplementary Fig. 1B), and the average gaze positions were calculated 50 ms before and after the saccade to determine whether they were near or inside the regions. The distance to a feature was defined as the minimum Euclidean distance between the gaze position and any point on the manually drawn contour line of the feature. (2) The amplitude of the saccade was greater than 2°. This second criterion ensured that the saccade was not small enough to occur right around the boundary of the two features but was large enough to cause a considerable change in the retinal input. The number was determined through the manual inspection of saccade patterns and the distribution of saccade amplitudes (Fig. 2E). (3) The saccade started at least 50ms after the participant fixated on the stimulus and at least 50ms before the participant’s response. This condition ensured that it occurred during decision formation.

To examine how stimulus fluctuations influenced the participants’ decisions depending on their gaze positions, we defined the fixated and unfixated features in each cycle of stimulus fluctuations in several analyses (Figs. 3, 4, Supplementary Fig. 8A, B). We first averaged eye positions within each of the 106.7 ms fluctuation cycles and then checked whether the averaged position was inside or near (< 1.5° for the face tasks and < 0.5° for the car task) the region of a feature circumscribed manually. If so, the feature was defined as fixated, whereas the other feature was defined as unfixated. This definition follows the criteria used to define cross-feature saccades above. If the average eye position was outside the range of both features, the fixated feature was not defined, and the corresponding cycle of stimulus fluctuations was excluded from the analysis. A fluctuation cycle was also excluded if there was a cross-feature saccade within this period.

During quantitative analyses and model fitting (Figs. 3G, 4, Supplementary Figs. 6, 7, 8C), we also calculated the distance between the participant’s gaze position and each object feature each time in a trial. This distance followed the definition described above as the minimum Euclidean distance between the gaze position and any point on the contour line circumscribing the region of the informative feature (Supplementary Fig. 1B). If the gaze position was within the circumscribed region, the distance was set to zero. Thus, this definition is agnostic of where the exact center of the informative features is.

#### Psychometric and chronometric functions

To quantify behavioral performance in the free saccade task, we fitted the following logistic function to the choice data of each participant for each stimulus condition (Fig. 1C top):

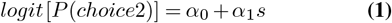

where *logit*(*p*) = *log*(*p/*1 − *p*), s is the nominal stimulus strength of a trial ranging from –1 (–100% morph level) to +1 (+100% morph level), and *α*_*i*_ are regression coefficients. *α*_0_ quantifies the choice bias and *α*_1_ quantifies the slope of the psychometric function.

The relationship between stimulus strength and the participants’ mean reaction times (RTs) was assessed using a hyperbolic tangent function (Fig. 1C bottom):

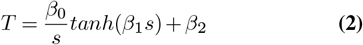

where T is the mean RTs in seconds and *β*_*i*_ indicates the model parameters. *β*_0_ and *β*_1_ determine the stimulusdependent changes in RTs, whereas *β*_2_ quantifies the portion of RTs independent of the stimulus strength.

Behavioral performance in the guided saccade task (Fig. 5) was quantified using the following logistic function

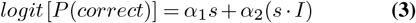

where an indicator variable, *I*, was used to quantify the difference in the slope of the psychometric functions between two conditions. For example, when we compared behavioral performance between the “Both” and “Pre only/Post only” conditions in the guided saccade task (Fig. 5B, E), *I* was set to 1 in the former condition and 0 in the latter condition. We fitted the above function to individual participants’ data and examined the performance difference between conditions by testing if *α*_2_ was significantly different from zero using *t*-test across participants. The function did not have a bias term because it was fit to the probability of correct, which is 0.5 at zero stimulus strength by definition.

#### Joint psychometric functions of features across saccades

To directly test whether participants used fixated features before and after cross-feature saccades, we plotted their choice performance as a function of the morph levels of these features (Fig. 3A-C). For example, if a participant first fixated on the eye region and then made a saccade to the mouth region before committing to a choice, we computed the average morph fluctuations in the eye region before the saccade as well as the average in the mouth region after the saccade (Fig. 3A). We then projected each trial in a 2D space defined by the morph levels before and after a saccade. In this space, we computed the probability of choice of the trials in a Gaussian window with a standard deviation of 5% and visualized the probability of choice by drawing iso-probability contours at 10% intervals (Fig. 3B). Similarly, if a participant made two cross-feature saccades, we plotted their performance as a function of the first, second, and third fixation features (Fig. 3C). Since the participants made fewer than three cross-feature saccades in most trials (Fig. 2F), we did not consider trials with more saccades.

To test the significance of the influence of each fixated feature on participants’ choices, we performed the following logistic regression for trials with one cross-feature saccade:

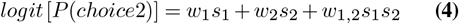

where *s*_1_ and *s*_2_ correspond to the morph levels of the features fixated the first and second times in a trial, respectively. *w*_1_ and *w*_2_ are linear coefficients, whereas *w*_1,2_ is a coefficient of the multiplicative interaction term. For trials with two cross-feature saccades, we used

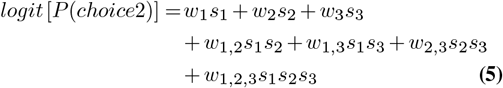

where *s*_1_, *s*_2_, and *s*_3_ corresponded to the morph levels of the features fixated first, second, and third times in a trial. We performed regression using all trials within individual participants and performed a two-tailed *t*-test across participants to test the significance of the contribution of each feature.

#### Psychophysical reverse correlation

To test whether features at different points in time across saccades affected the participants’ choices, we performed psychophysical reverse correlations (30, 35) (Figs. 2H, 3E, 4, Supplementary Figs. 5-9). Psychophysical kernels (*K*_*f*_ (*t*)) were calculated as the difference in the average fluctuations of morph levels conditional on the participant’s choices:

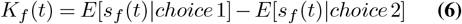

where *s*_*f*_ (*t*) represents the morph level of feature *f* at time *t*. This analysis only used trials with low stimulus strength (nominal morph level, 0-12%). For the non-zero strength trials, the mean strength was subtracted from the fluctuations, while the residuals were used for the reverse correlation. In Figure 2H, we averaged the kernels for each feature over time when the participants viewed the feature. For the guided saccade tasks (Fig. 5D, G), we averaged the two cycles of stimulus fluctuations both before and after the saccade to calculate the kernels (but see Supplementary Fig. 9G, H for unaveraged kernels).

When plotting the time course of psychophysical kernels (Figs. 3E, 4, Supplementary Figs. 5-7), we sorted individual stimulus fluctuations depending on which feature the participant fixated on during that cycle of fluctuations, and then generated kernels for the fixated and unfixated features. When a participant’s gaze position could not be classified into a feature or a cross-feature saccade occurred during a cycle of fluctuation, it was excluded from the analysis. Since the stimulus fluctuation started when participants fixated on an image (see above), the kernels aligned to stimulus onset started from the moment of fixation and were calculated up to the first cross-feature saccade or up to 1 s in the trials without saccades. The kernels aligned to the participants’ responses were calculated using stimulus fluctuations after the last cross-feature saccade or using the fluctuations for 1 s from the response when there was no saccade in a trial. For the kernels aligned to saccades, we used five stimulus cycles before and after the saccade onset. Figure 4D shows an example trial with only one saccade, but if there were more than one cross-feature saccade, all were used when computing the kernels. For the kernels shown in Figures 3E and 4, we averaged the kernels across all participants. The kernels of the individual participants can be found in Supplementary Figure 6. Three-point boxcar smoothing was applied to the temporal kernels for denoising. However, we did not perform smoothing when evaluating the fitting quality (*R*^2^).

To further quantify how gaze position modulated the contribution of local features to decisions, we realigned the same stimulus fluctuations according to the distance between the participants’ gaze position and each feature location (Fig. 3F, G). As explained in the “Detection of cross-feature saccades and quantification of saccade patterns” section above, we averaged the gaze positions during each cycle of stimulus fluctuation and computed its distance from any point on the counter line circumscribing the region of each feature (Supplementary Fig. 1B). If the gaze position was within the circumscribed region, the distance was set to zero. If a saccade occurred during one cycle of stimulus fluctuation, the fluctuation was excluded from the analysis. We then sorted the fluctuations according to the calculated distance and generated psychophysical kernels at each distance *d* as

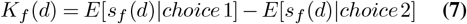

where *s*_*f*_ (*d*) is the morph fluctuation of feature *f* at distance *d*. This was calculated using stimulus cycles concatenated across the trials with low stimulus strength (nominal morph level, 0-12%) within individual participants.

#### Calculation of saccade frequency and probability

We calculated the frequency of participants’ saccades for each stimulus strength to test any potential dependence on stimulus difficulty (Fig. 6B). Frequencies could not be simply estimated by dividing saccade counts by trial duration (i.e., RTs) because saccades tended to be periodic. Suppose that saccades occur every 400 ms regardless of stimulus strength. If the average RT was 500 ms for one stimulus and 700 ms for another, saccade counts are expected to be one per trial, and thus, saccade counts per time tend to be underestimated for stimuli with longer RTs. Therefore, we had to match the RT distributions across stimulus strengths for a proper comparison. We generated RT histograms with 100 ms intervals and performed histogram matching by randomly subsampling trials from each stimulus strength. We then counted the total number of saccades in these subsampled trials and divided this number by the total stimulus duration across the trials. The frequencies were calculated in this manner for individual participants and then averaged (Fig. 6B).

We further took an alternative approach to estimate saccade frequency without matching RT distributions (Fig. 6C). In this method, we calculated the number of saccades that occurred at each time point (100 ms bins) and divided it by the number of trials whose RT was longer than that time point. This saccade probability is not affected by the interaction of RTs and saccade timing outlined above because this metric does not depend on the duration of trials after each time point to compute the probability. However, the results can be noisy, particularly for later time points, because fewer trials contribute to the calculation. We therefore classified trials into two groups (easy: > 20% morph level, difficult: < 20%) to perform this analysis.

### Model fit and evaluation

To quantitatively examine whether the participants’ choice behavior during the free-saccade task could be explained by the integration of sensory evidence over saccades, we constructed a simple extension of evidence accumulation models widely used to explain behavioral data in a variety of perceptual decision-making tasks (28, 29). We also developed multiple alternative models to confirm that no alternative mechanisms account for the behavioral data. In what follows, we first describe the expression and fitting procedure for the main model and then extend them to the alternative models.

#### Main model

Our main model is an extension of the drift-diffusion model, which considers multiple informative image features and their distances from the participants’ gaze positions (Fig. 4A). The model was extended from that of a previous study that was demonstrated to accurately explain human face categorization behavior measured under stable fixation conditions (29). This previous model first linearly integrates the fluctuations of the local features (*s*_*i*_(*t*) for feature *i* at time t):

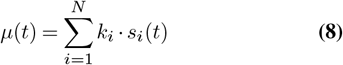

where *µ* is momentary evidence for the model, *N* is the number of features, and *k*_*i*_ is the sensitivity parameter for each feature *i*. Momentary evidence is then accumulated over time to form the decision variable (*v*) at each time *t* as

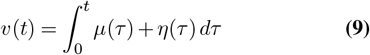

where *η*(*τ*) represents internal (neural) noise in the sensory, inference, or integration processes, assumed to follow a Gaussian distribution with mean 0 and SD *σ*(*t*). When the decision variable (*v*(*t*)) reaches an upper or lower bound (+B or -B), the model commits to a decision associated with the bound. Reaction time was defined as the time required to reach a bound plus a non-decision time including sensory and motor delays. The non-decision time was drawn from a Gaussian distribution with a mean of *T*_0_ and an SD of 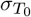.

Our present model extends the above formalism by incorporating participants’ gaze positions as a factor that influences the informativeness of local features (39, 74, 75). We added one free parameter (λ) that quantified the degree to which the informativeness of each feature decays as a function of the distance from the gaze position (i.e., visual eccentricity). The decay was expressed as an exponential function based on the previous studies that successfully modeled visual acuity as a function of visual eccentricity (8, 40). We also tested a linear decay function and confirmed that it yielded similar results (Supplementary Fig. 7B). In the exponential model, Eq. 8 was modified as:

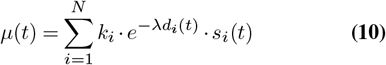

where *d*_*i*_(*t*) is the Euclidean distance (in units of visual angle) between the gaze position and object feature *i* at time *t*. As mentioned above, the distance was defined as the minimum length between the gaze position and any point on the counter line manually drawn to circumscribe each feature (Supplementary Fig. 1B). If the gaze position was inside the circumscribed region, the distance was set to zero. During the saccades, both momentary evidence and diffusion noise were set to zero to simulate the absence of visual input.

Once the momentary and accumulated evidence is defined as above, we can numerically derive the probability that the decision variable has value *v* at time *t* by solving the Fokker-Planck equation:

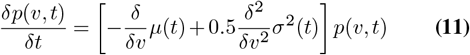

where *p*(*v, t*) denotes the probability density. The accumulation process started from zero evidence and continued until the decision variable reached one of the two bounds (±*B*), indicating two choices. Thus, the partial differential equation above has the following initial and boundary conditions:

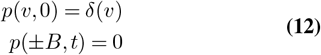

where *δ*(*v*) denotes the Dirac delta function. The diffusion noise (*σ*(*t*)) was set to 1, and the bound and drift rate were defined in a unit of diffusion noise. The RT distribution for each choice was obtained by convolving the distribution of bound crossing times with the distribution of non-decision time (a Gaussian distribution with a mean of *T*_0_ and an SD of 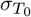). The SD, 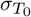, was always set to one-third of *T*_0_ to reduce the number of the free parameters.

Overall, our main model had five degrees of freedom: decision bound height (*B*), sensitivity parameters for two features (*k*_1_, *k*_2_), mean non-decision time *T* 0, and the decay rate of visual sensitivity λ. We fit the model parameters by maximizing the likelihood of the joint distribution of the observed choices and RT distributions of individual participants in each stimulus condition (30). Given a set of parameters, the stimulus fluctuations and participant’s gaze points in each trial were used to calculate the RT distributions of the two choices according to the model formulation above. These distributions were used to calculate the log-likelihood of the observed choice and RT for individual trials. These log-likelihoods were summed across the trials to calculate the likelihood function for the dataset. We used a simplex search method (*fminsearch* in Matlab) to determine the parameter set that maximized the summed likelihood. To avoid local maxima, we repeated the fitting process from multiple initial parameter sets and selected the set that converged to the largest likelihood as the final result. Because maximum likelihood estimation is sensitive to outliers, we excluded trials with reaction times greater than three SDs from the mean for each stimulus strength during model fitting. Fitting was performed for each participant and included the trials with all stimulus strengths. The fitting performance was quantified using the coefficient of determination (*R*^2^) for the joint distributions of choices and RTs. For each morph level, we generated the RT distribution for each choice (bin size, 100 ms) and computed the *R*^2^ between the data and model outputs after concatenating the bins for all morph levels and choices. The fitting curves shown in Fig. 4B-D are the averages across participants.

#### Alternative models

To examine whether different mechanisms accounted for the behavioral data, we developed multiple alternative models. They included the “gaze independent” model, which had constant sensitivity to each local feature regardless of the participant’s gaze position, the “evidence reset” model, which resets the accumulated evidence every time the participant makes a cross-feature saccade, and the “independent accumulator” model, which does not integrate evidence across saccades but accumulates the evidence from two features independently. These models were fitted to the behavioral data using the abovementioned procedure.

The gaze-independent model was designed to test whether the information of participants’ gaze positions was necessary to account for their behavior. In our main model, the sensitivity to each feature was modulated according to the distance between the gaze position and the feature (Eq. 10), whereas the gaze-independent model removed this term and computed momentary evidence assuming that the sensitivity to each feature is constant regardless of gaze position (thus using Eq. 8) to determine the drift rate. The other components of the model were the same as those used in our main model. This model has one fewer parameter (4) than our main model.

The evidence-reset model was created to test the possibility that, when sensory evidence from one feature was insufficient to form a decision, people would make a saccade to the other feature and restart their decision-making process. To simulate this, the model resets the accumulated evidence to zero after a cross-feature saccade. Thus, the choices and RTs of the model were based solely on the feature fixated on after the last cross-feature saccade in a trial. To fit this model, we extracted the timing of the last cross-feature saccade and the feature fixated afterward from each trial and simulated the bounded evidence accumulation using them to predict the choice and RT of that trial. The model was equivalent to our main model if a trial did not contain a cross-feature saccade. The model becomes unrealistic when the last cross-feature saccade was too close to the RT of a trial; we thus did not count saccades that occurred within the mean non-decision time plus one standard deviation of non-decision time (i.e., 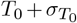) from the RT. Besides this resetting mechanism, all components of our main model, including the sensitivity of each feature and the dependency of sensitivity on the gaze positions, were preserved in this model. The number of parameters in this model is the same as that in our main model.

The independent-accumulator model tested the possibility that the evidence was not integrated across saccades. Instead, it independently accumulated evidence for the features fixated on before and after saccades and committed to a choice based on evidence from one of them. To explain the RTs, the model computed the timing of reaching a decision bound using the accumulated evidence of the last fixated feature, as in the evidence reset model. However, the choice was determined by the total evidence accumulated for one of the features during the entire stimulus presentation period when that feature was fixated on. We considered two mechanisms to determine the features used to make a choice. One mechanism involved random selection; with a probability *η* (a free parameter to be fit), the feature lastly fixated was used to determine a choice, and with a probability 1 − *η*, the other feature was used. The second involves the selection of features with greater evidence. After reaching a bound, the model compared the total evidence accumulated for each feature and used the one with the greater absolute value to determine the choice.

The independent-accumulator model was fitted using the participants’ eye-movement patterns in each trial. We classified each period of fixation between cross-feature saccades into one of the two features and then calculated the sum of evidence across the periods for each feature. The model with the random selection mechanism had six parameters. The model that used the feature with higher evidence had five parameters.

To test fit performance, we computed the difference in the Bayesian information criterion (ΔBIC) between the main model and each of the alternative models (Fig. 4, Supplementary Fig. 7; positive values indicate poorer fits of the alternative models). When comparing the main model and the gaze-independent model, we summed the log-likelihood of all trials and averaged the sum across participants to derive the BIC. When comparing with the evidence-reset and independent-accumulator models, we found that the same procedure was not a fair comparison because these models required the timing of the last cross-feature saccade and thus had access to additional information for fitting choices and RTs. Therefore, we shuffled the model-predicted choice and RT distributions across trials within each morph level before computing the log-likelihood. This shuffling, which was also done for the main model, removed the additional information of saccade timing. However, these models predicted a log-likelihood of negative infinity for RTs before the last saccade. To address this, we reshuffled until there were no trials with infinite log-likelihood.

#### Generation of model psychophysical kernels and RT distributions

The models above were fit to the choices and RTs, but the model formulation does not prescribe its psychophysical kernel. Therefore, we relied on simulations to estimate the model kernels. We created 10^5^ simulated trials with stimulus strengths ranging from 0-12% using the same stimulus distributions as in the main task (i.e., Gaussian distribution with 20% SD). The model responses for these trials were simulated using the same parameters fitted for each participant. We then used the simulated choices and RTs to calculate model psychophysical kernels, following the same procedure used for the human data (Fig. 4D, F, H, J, Supplementary Figs. 6, 7). Thus, the model kernels were not directly fitted to the participants’ kernels but were generated from an independent set of stimulus fluctuations, making the comparison of data and models informative. Similarly, the RT distributions of the models (Fig. 4C) were generated using simulations with an independent set of morph fluctuations to ensure an accurate comparison of the data and models.

To generate the model predictions, it was necessary to simulate eye movement data because the models needed to compute the distance between gaze positions and feature locations to calculate the strength of momentary evidence (Eq. 10). To generate realistic eye data, we used the participants’ actual eye movement data from randomly selected trials; however, when the duration was shorter than the duration required for model simulations, we extended the eye data in two different ways. For the main model, we stitched a chunk of the eye trace obtained from another trial such that it could be smoothly connected to the end of the eye data. To do so, we looked for a chunk that started at a position less than 0.3° distance from the endpoint of the eye data. We repeated this stitching procedure until the eye trace reached the desired length. For the evidence reset and independent accumulator models, we simply extended the last eye position to become the desired length of the eye trace because the model had to assume that no saccade occurred during this extended period.

### Ideal observer analysis for the guided saccade task

In the guided saccade task (Fig. 5), we examined whether the participants’ performance could be accounted for by the optimal integration of the evidence before and after a saccade (Supplementary Fig. 9). The task had “Pre only” and “Post only” conditions where a stimulus was only shown before or after a saccade, and “Both” condition where a stimulus was shown in both epochs (Fig. 5B). To build an ideal observer model, we first estimated the precision of the participant’s judgment of the stimulus in the “Pre only”, “Post only”, and “Both” conditions 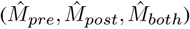 assuming Gaussian judgment noise:

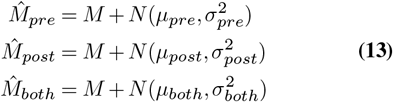

where *M* is the actual stimulus value (morph level), *µ*_*pre*_, *µ*_*post*_, and *µ*_both_ are biases in the judgment,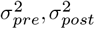and 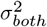 are variances in the judgment, and *N* represents the Gaussian distribution. These biases and variances could be estimated by fitting a cumulative Gaussian distribution to the psychometric function of the “Pre only”, “Post only”, and “Both” conditions, respectively.

An ideal observer model that optimally combines evidence from “Pre” and “Post” epochs makes the following maximum a posteriori estimate (14, 76, 77):

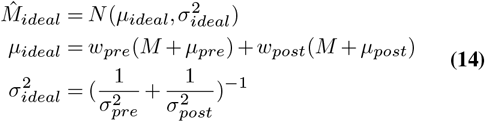

where

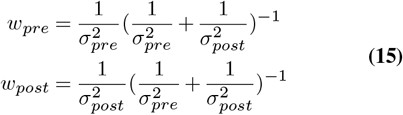

The obtained 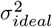 corresponds to the variance of the judgment by the ideal observer. We compared this value against 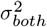 calculated above to test the optimality of the integration (Supplementary Fig. 9A-B). We further performed the same analysis for the no-saccade condition (Supplementary Fig. 9C-D).

## Acknowledgements

We thank Tianlin Luo for comments on earlier versions of the manuscript. This work was supported by the National Science and Technology Innovation 2030 Major Program (Grant No. 2021ZD0203703, G.O.), Strategic Priority Research Program of the Chinese Academy of Sciences (XDB1010202, G.O.), National Natural Science Foundation of China (Grant No. 32371077, G.O.), and the National Natural Science Fund for Excellent Young Scientists Fund Program (overseas).

## Data and code availability

Data and code used in this study will be deposited on the Open Science Framework (OSF) website upon official publication.

## Supplementary figures

**Fig S1.**
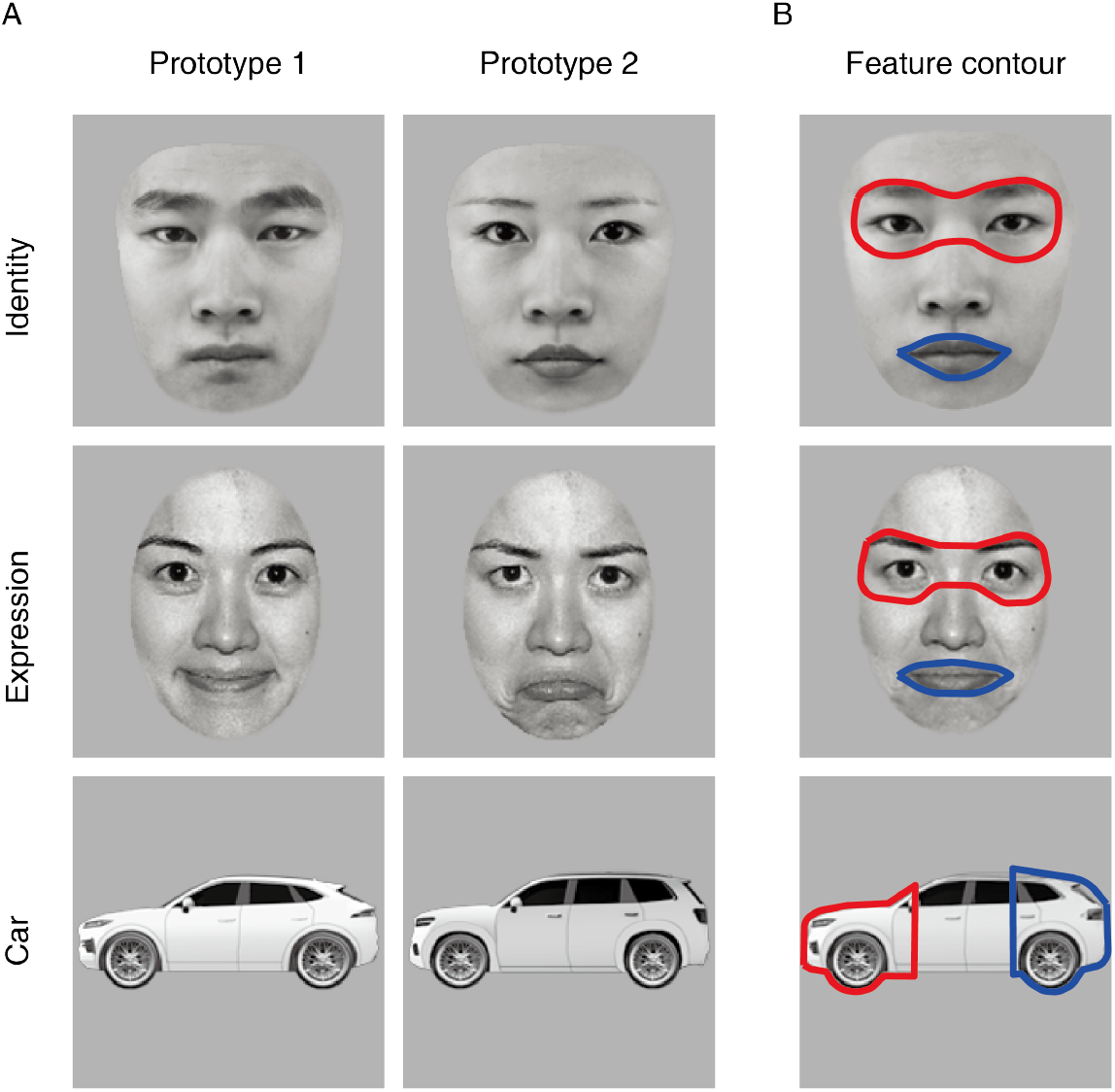
Prototype images and definitions of feature regions. (**A**) The two prototype images used in each of the three stimulus conditions. The face images were from the Nim Face set (33) and the Tsinghua Facial Expression Database (34) and presented with permission. The same face images were used in the subsequent figures. (**B**) The manually defined regions of the two features used to determine which feature the participants fixated on at each moment (red: feature 1, blue: feature 2). We calculated the minimum Euclidean distance between a gaze position and any point on the contour line and considered that the participant looked at the feature if the distance was smaller than 1.5° for the face tasks or 0.5° for the car task. If the gaze position was inside the circumscribed region, the distance was defined as zero. These numbers were chosen according to the manual inspection of participants’ overall gaze positions.

**Fig S2.**
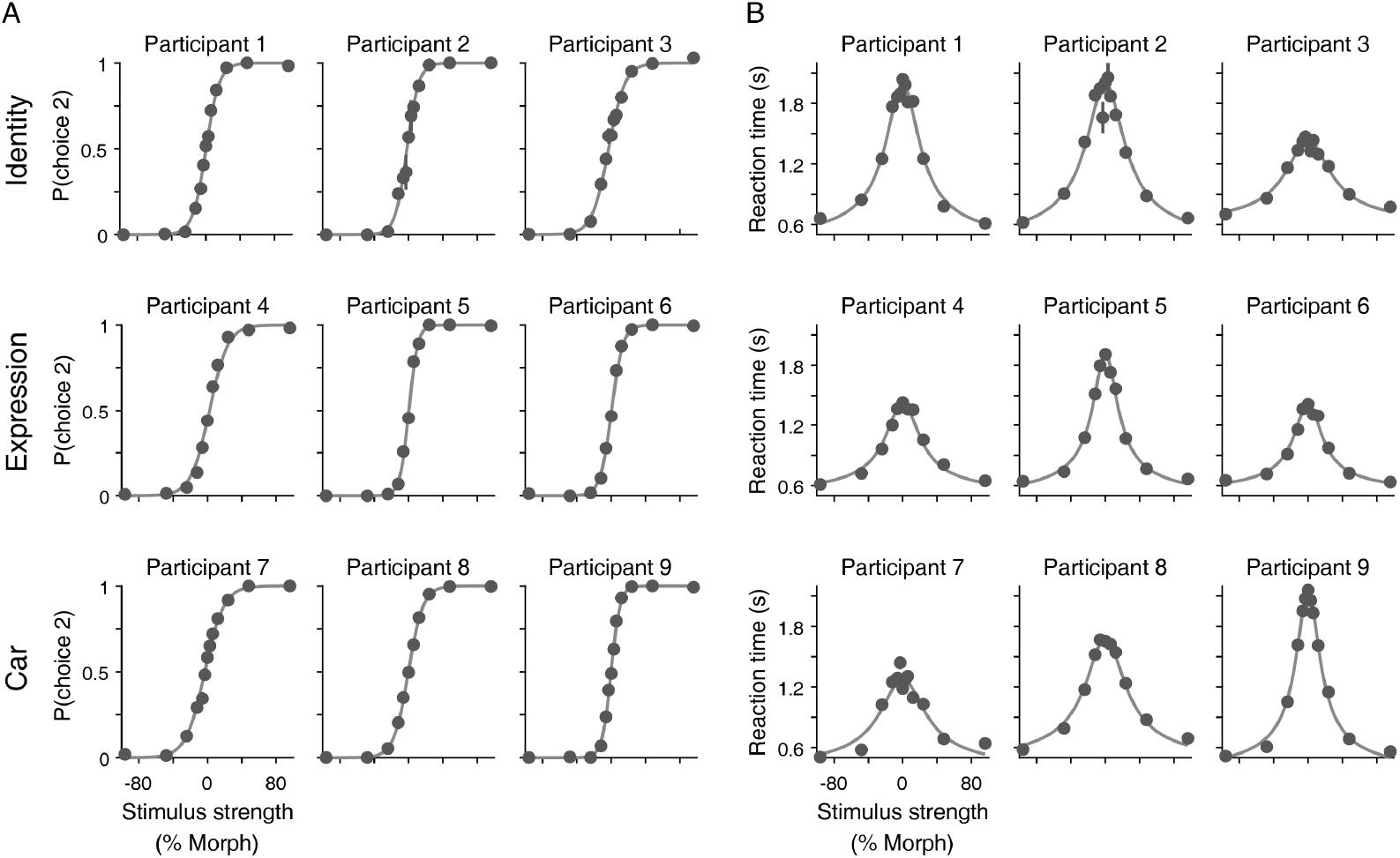
Psychometric and chronometric functions for individual participants. Individual participants’ plots for Fig. 1C. Participants showed stereotypical choice accuracy (**A**) and mean reaction times (**B**) as a function of the mean morph levels. Lines represent the logistic and hyperbolic tangent fits for psychometric and chronometric functions, respectively.

**Fig S3.**
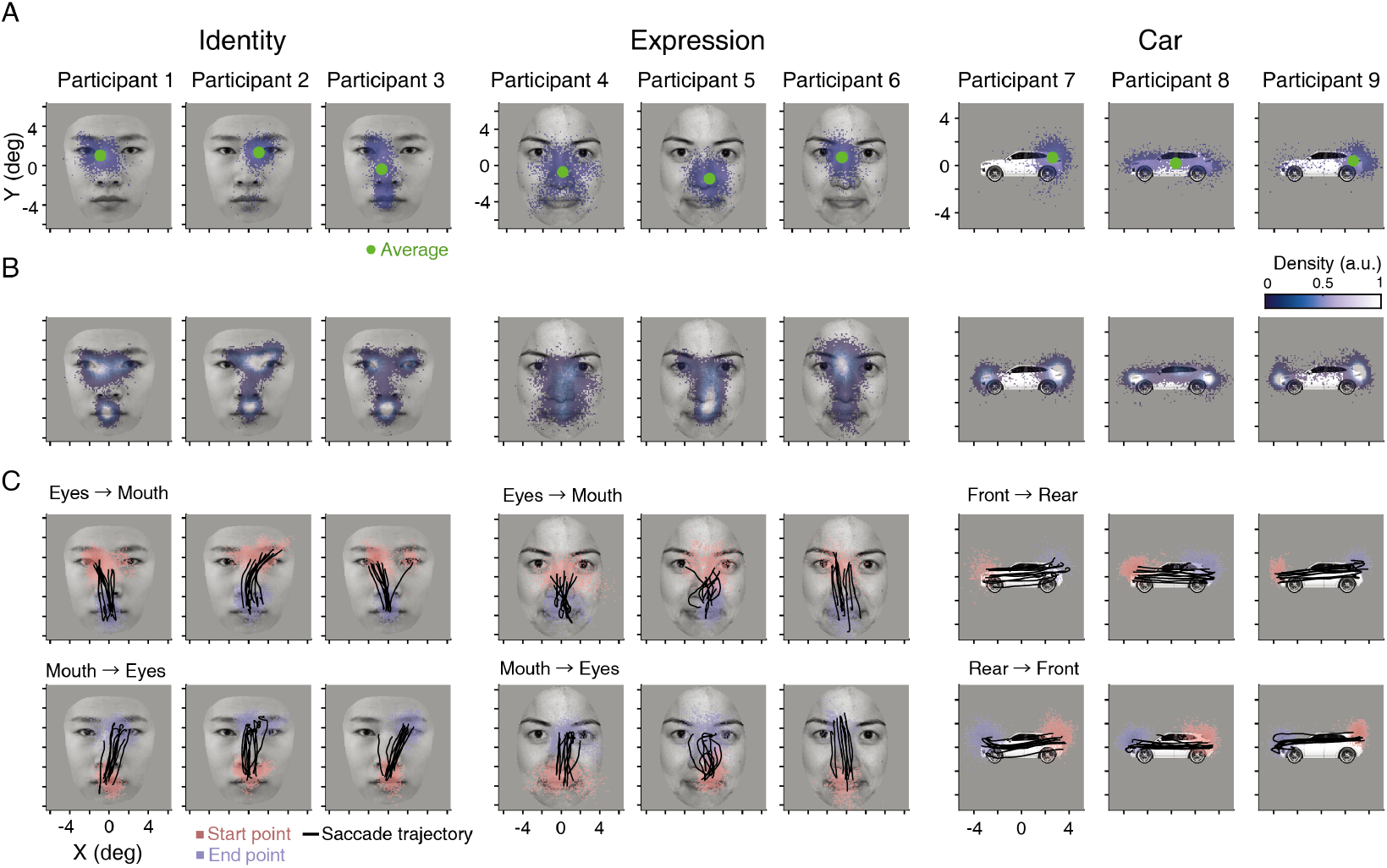
Fixation and saccade patterns were largely consistent across participants. Individual participants’ plots for Fig. 2A-C. (**A**) Scatter plots of the initial fixation positions after stimulus onset for each participant. Blue dots represent individual trials, while green dots represent the average. (**B**) Density plots of the fixated positions during the entire stimulus viewing period. (**C**) Example saccades spanning the two informative features.

**Fig S4.**
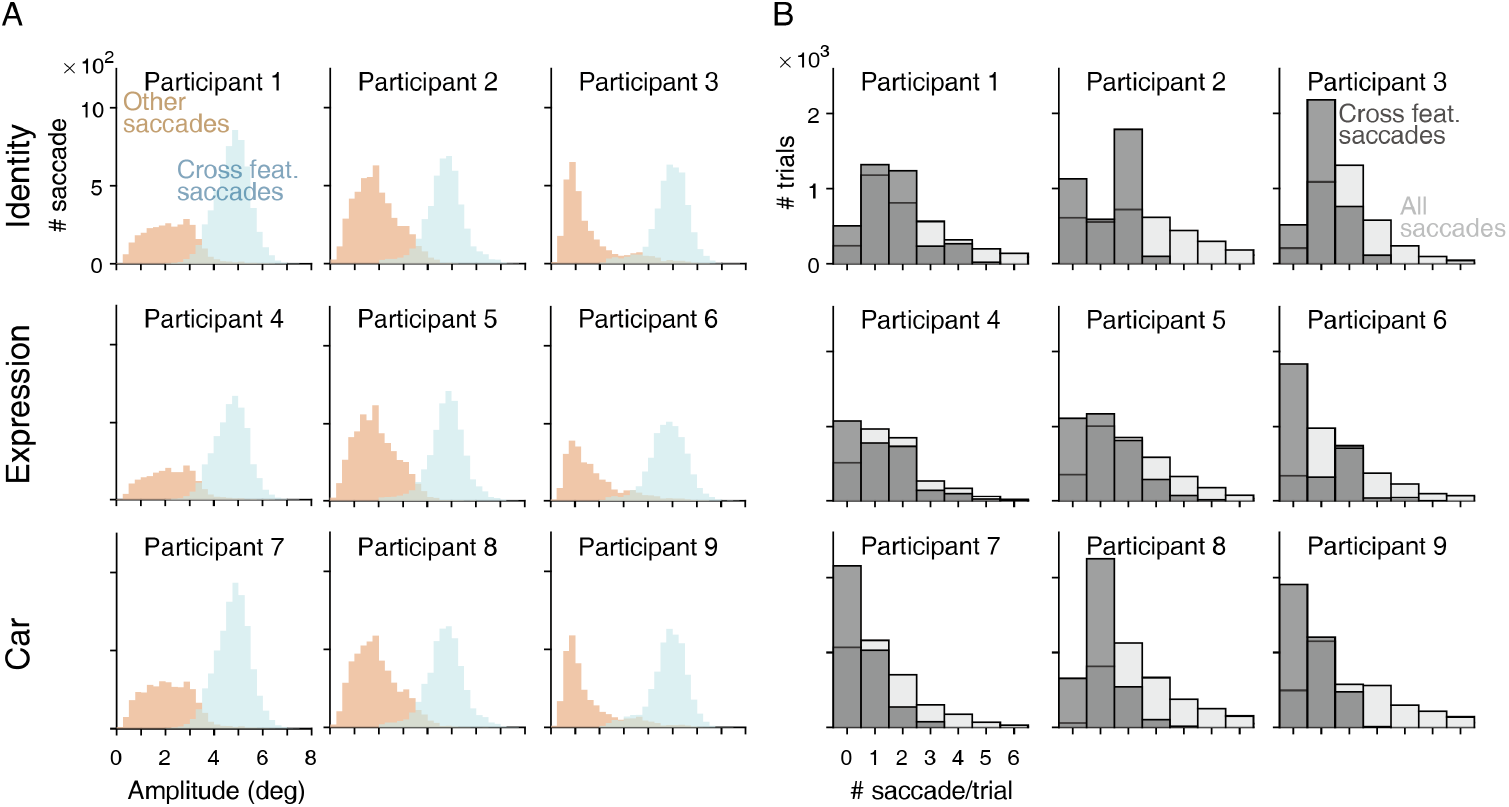
The amplitude and frequency of saccades were largely consistent across participants. Individual participants’ plots for Fig. 2D-F. (**A**) The amplitude distributions of cross-features and other saccades for each participant. (**B**) The distributions of saccade counts per trial for each participant.

**Fig S5.**
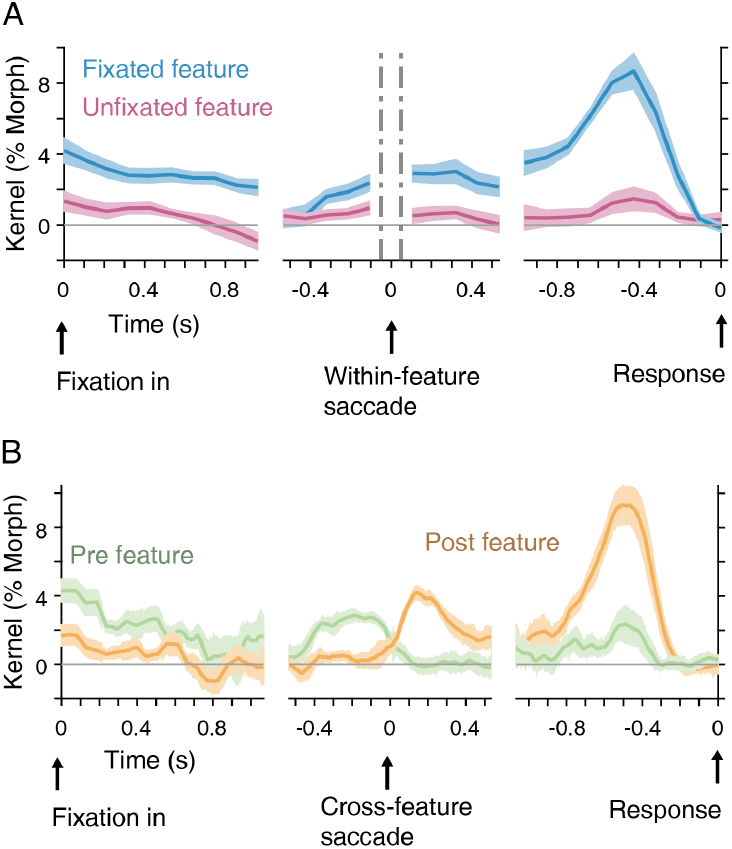
Psychophysical kernels plotted in different forms. (**A**) When psychophysical kernels were plotted around the time of the within-feature saccades, they showed the continuous weighting of the same, fixated features across saccades. This supports the integration of evidence across small saccades and complements the results for cross-feature saccades (Fig. 3E). Shaded regions indicate S.E.M. across participants. (**B**) Psychophysical kernels (Fig. 3E) plotted with a higher temporal resolution. In the experiments, morph levels fluctuated every 106.7 ms (Fig. 1B), and the main figures (Figs. 3E, 4) were generated by maintaining this resolution. However, saccades could occur at any time during the cycles of morph fluctuations. In the main figures, the offsets between morph fluctuations and saccades were rounded to generate kernels; however, this panel plotted kernels after recasting morph fluctuations at 1 ms resolution assuming that a morph level is constant during a 106.7 ms cycle. The kernels were smoothed with a 50 ms boxcar function. The results revealed a slight decrease in the amplitude of pre-saccade feature before saccades and an increase in the amplitude of post-saccade features after saccades over 100-200 ms (see Wolf and Schütz (2015) for comparison). However, this should be interpreted with caution, as the data are limited by the temporal resolution of the experimental paradigm.

**Fig S6.**
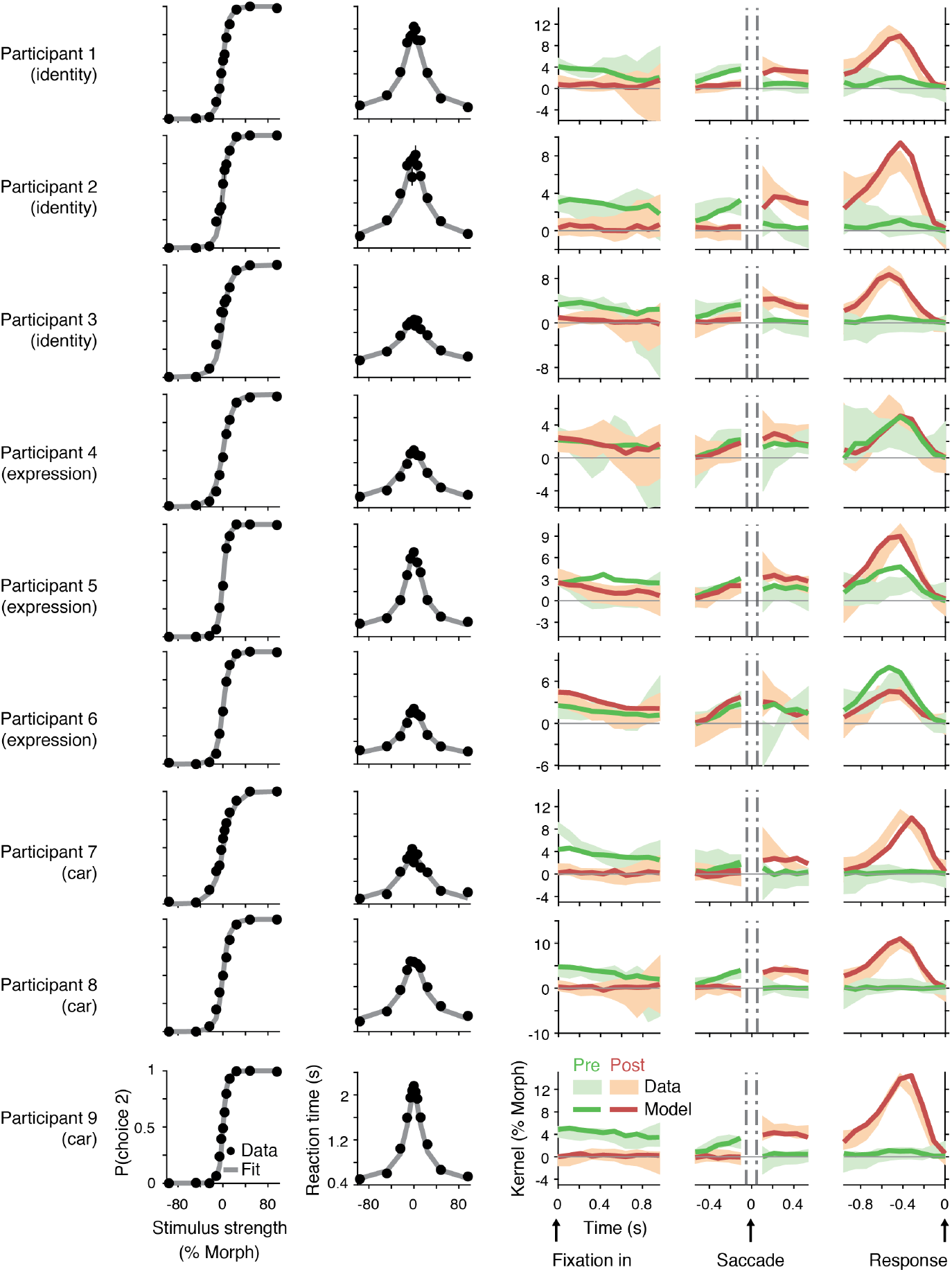
The main model accounts for choice, reaction times, and psychophysical kernels of individual participants. The main model (shown in Fig. 4A) was fitted to individual participants’ data. The conventions are the same as for Figs. 4B and D. Participant 6’s data and model fit showed a larger kernel amplitude for the “Pre” feature in the post-saccadic period. This occurred because the participant had a high sensitivity to the mouth and tended to look at the mouth first (thus it tended to be the “Pre” feature), resulting in a larger kernel even after saccades.

**Fig S7.**
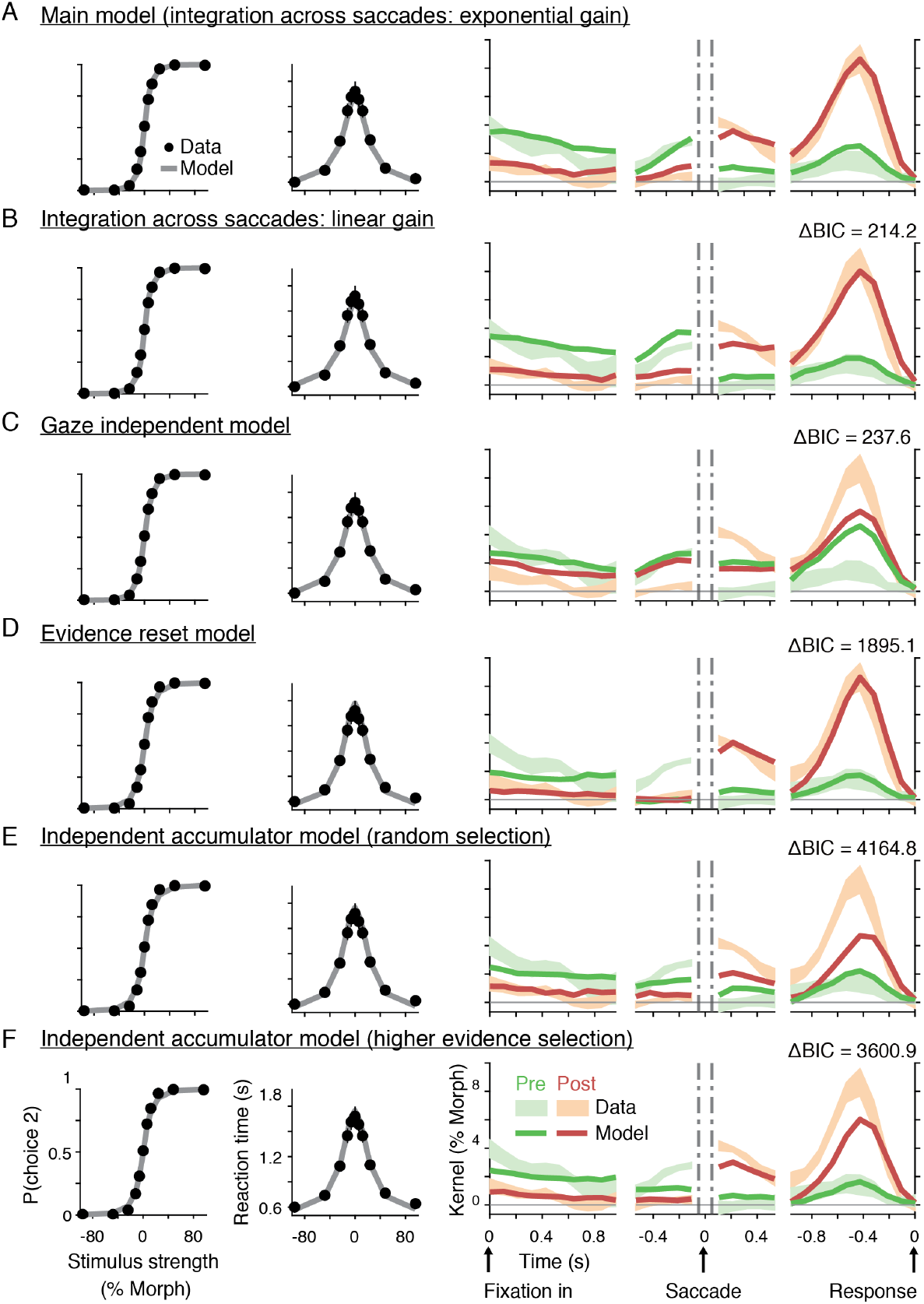
Comparison of psychometric and chronometric functions and psychophysical kernels across models. While alternative models could also fit the participants’ psychometric (left) and chronometric (middle) functions, only the main model (A) and its variant (B) accounted for the psychophysical kernels (right). Δ*BIC* indicates the difference in fit performance relative to the main model (positive values indicate poorer fits). (**A**) The plots for the main model (Fig. 4A) are the same as those for Fig. 4B and D. (**B**) The main model used an exponential gain function to explain the weighting of sensory features as a function of visual eccentricity (Fig. 4A inset), but changing it to a linear function yielded similar but slightly worse fitting. (**C**) The gaze-independent model (Fig. 4E, F). (**D**) Evidence reset model (Fig. 4G, H). (**E**) The independent-accumulator model (Fig. 4I, J), where the evidence from two features was randomly selected to make a decision. (**F**) Another variant of the independent-accumulator model, where the feature with higher evidence was selected to make a decision.

**Fig S8.**
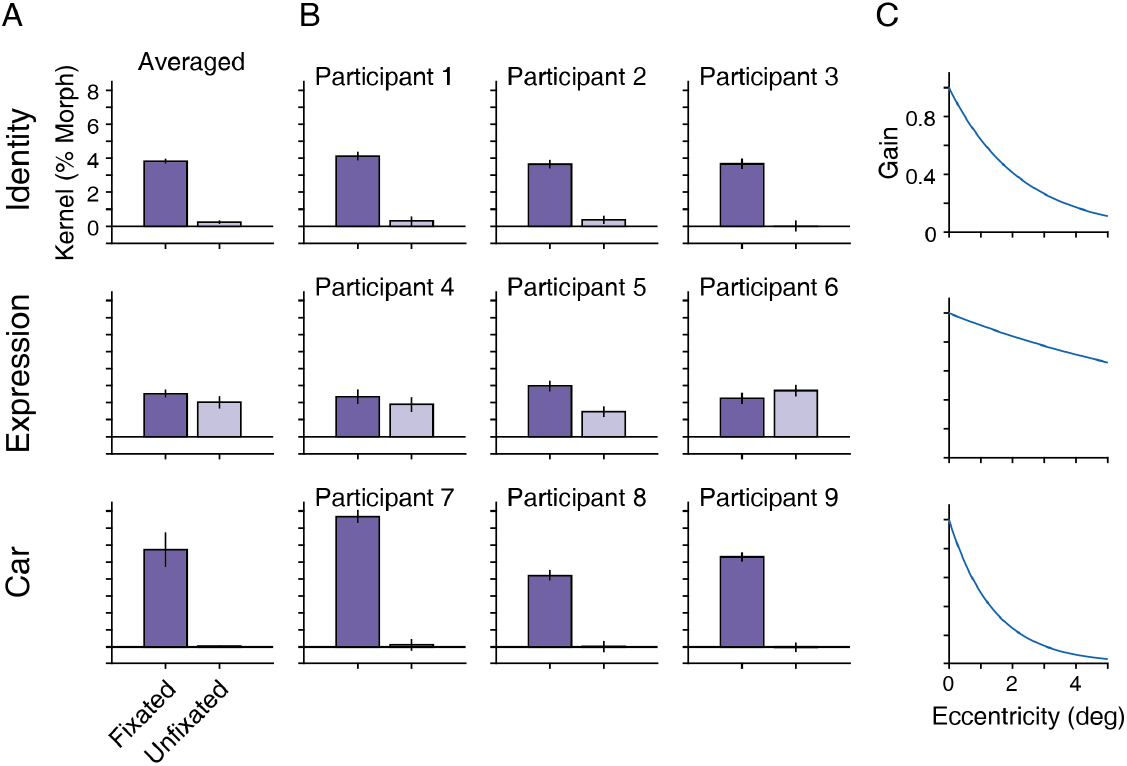
The reliance on the unfixated feature depended on the stimulus conditions. (**A**) We performed psychophysical reverse correlation (Eq. 6) after sorting the two informative features into fixated and unfixated ones at each stimulus fluctuation cycle. The panels show the average of the three participants in each of the three stimulus conditions shown in Fig. 1A. In the identity and car conditions, the kernel of the fixated feature was large, while the kernel of the unfixated feature was near zero. In the expression condition, in contrast, the kernels of the fixated and unfixated features showed similar amplitudes. This is consistent with the psychophysical kernels aligned to the distance from the gaze position (Fig. 3G), which revealed a sharp dependency on the distance in the identity and car condition and a flat amplitude over a range of distances in the expression condition. (**B**) These results were consistently observed in individual participants. Although these results remain preliminary due to the limited number of participants, they nevertheless indicate that the extent of spatial integration is task-dependent. (**C**) Visualization of the gaze-dependent gain in our model (Fig. 4A, inset) based on the fitted parameter (λ) also supports broader spatial sampling in the expression condition. The parameter λ was fitted for individual participants, while the plots are based on λ averaged across them for each stimulus condition. The gain at zero-degree distance was always 1.

**Fig S9.**
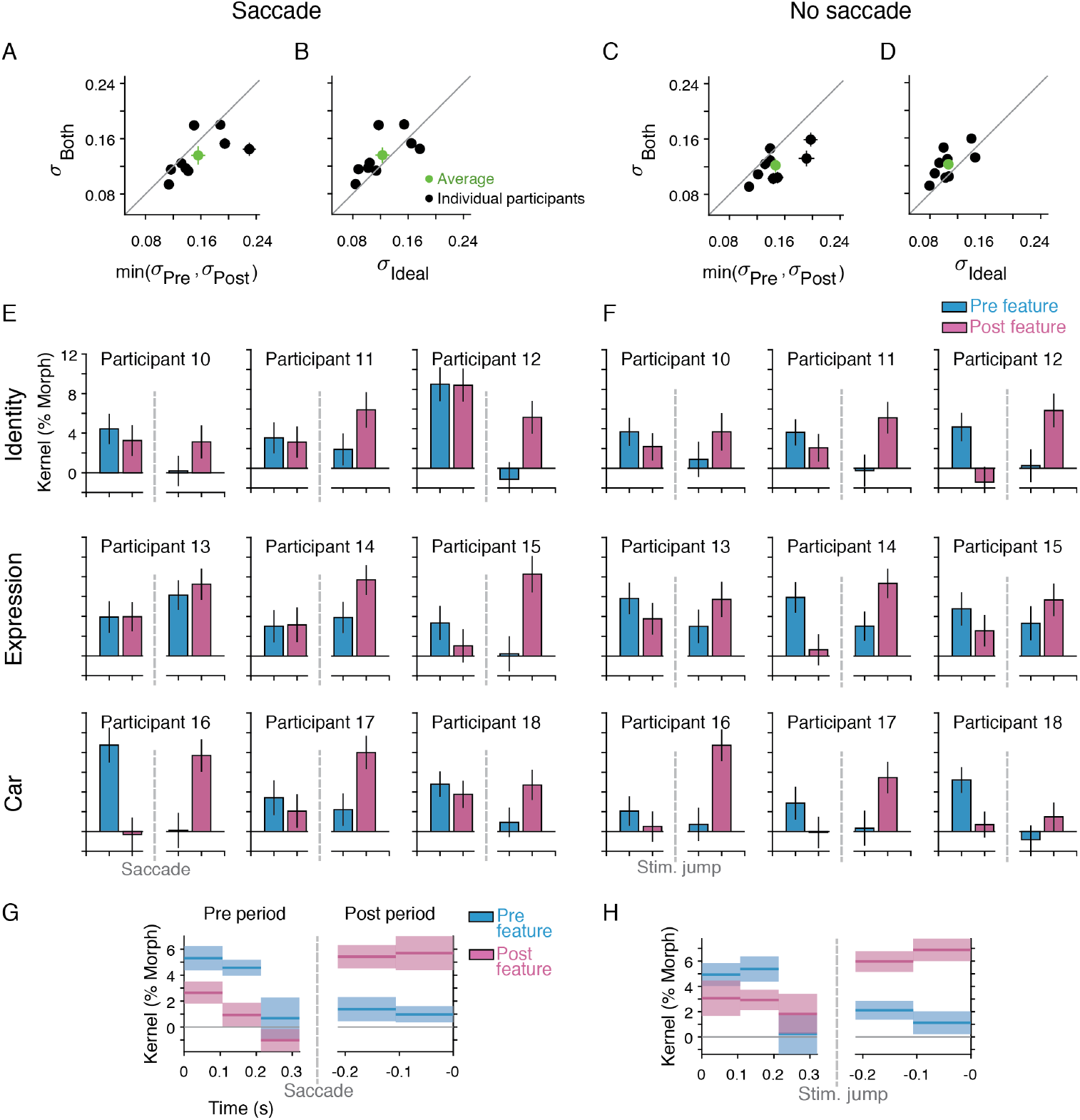
Near-optimal integration of features with and without saccades in the guided saccade task. (**A**) To examine the optimality of evidence integration in the guided saccade task (Fig. 5), we quantified the behavioral accuracy for the “Both”, “Pre only”, and “Post only” conditions (Fig. 5B, E) using Eq. 13, which provides us the standard deviation (SD) in the participants’ estimate of a stimulus. If participants integrate evidence, the SD in the “Both” condition should be smaller than the minimum SD between the “Pre only” and “Post only” conditions. In support of this, most participants showed smaller SDs in the “Both” condition (*t*_(8)_ = 1.93,*p* = 0.045, one-tailed *t*-test). Error bars indicate the S.E.M. (**B**) We further tested the optimality of this integration by constructing an ideal observer model (14, 76, 77) that optimally integrates evidence of the “Pre only”, and “Post only” conditions (see Methods; Eq. 14). The SDs of this ideal observer were similar to the participants’ SDs in the “Both” condition (*t*_(8)_ = 1.45,*p* = 0.093), although the participants’ performances looked slightly worse than those of the ideal observer. The result is consistent with near-optimal integration of Gabor orientations across saccades identified in a previous study (14). (**C, D**) We observed similar patterns of near-optimal integration in the no-saccade condition (Fig. 5E). Participants’ performance in the “Both” trials was better than the minimum of the “Pre only” and “Post only” trials (*t*_(8)_ = 3.37,*p* = 0.0049) but slightly worse than the ideal observer (*t*_(8)_ = 2.50,*p* = 0.018). (**E, F**) Individual participants’ plots for Fig. 5D and G indicate that participants used both features before and after a saccade/stimulus jump to make a decision. The participant IDs started from 10 for disambiguation because different participants were involved in this task and the free saccade task. (**G, H**) Figure 5D showed psychophysical kernels for pre- and post-saccade features calculated after averaging the morph fluctuations (2-3 cycles) in each epoch. Instead, the plots here show the kernels without averaging these cycles. Consistent with Fig. 5D and G, both the pre and post features had positive kernels and their amplitudes were stable over time, except for the third cycle of the pre period, where a saccade or stimulus jump often occurred early in this cycle.

**Fig S10.**
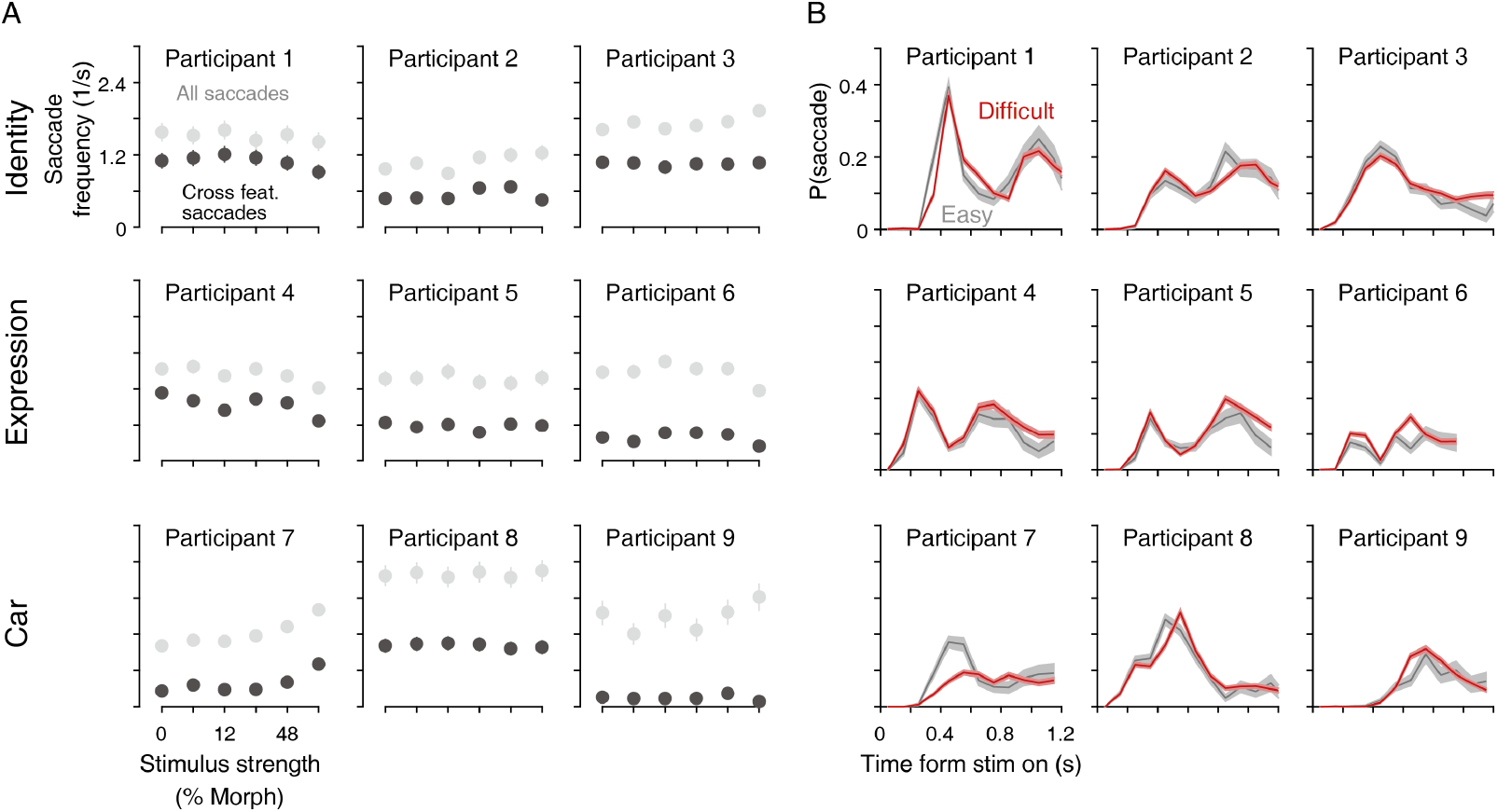
Eye movement patterns did not strongly depend on stimulus strength in the majority of participants. Individual participants’ plots for Fig. 6B, C. (**A**) The frequency of cross-feature saccades and all saccades was largely stable over a range of stimulus difficulties. As in Fig. 6B, the RT distributions were matched across stimulus strengths for proper comparison. Error bars indicate the S.E.M. across trials. (**B**) The probabilities of making cross-feature saccades at each moment were also largely similar between the easy (morph level: > 20%) and difficult (< 20%) trials, except for participant 7, who tended to make more saccades in easy trials. Error bars indicate the S.E.M. across trials.

